# BRAT1 - a new therapeutic target for glioblastoma

**DOI:** 10.1101/2024.07.08.602519

**Authors:** Alicia Haydo, Jennifer Schmidt, Alisha Crider, Tim Kögler, Johanna Ertl, Stephanie Hehlgans, Marina E. Hoffmann, Rajeshwari Rathore, Ömer Güllülü, Yecheng Wang, Xiangke Zhang, Christel Herold-Mende, Francesco Pampaloni, Irmgard Tegeder, Ivan Dikic, Mingji Dai, Franz Rödel, Donat Kögel, Benedikt Linder

**Affiliations:** Experimental Neurosurgery, Department of Neurosurgery, Neuroscience Center, Goethe University Hospital, 60528 Frankfurt am Main, Germany; Department of Radiotherapy and Oncology, Goethe University Hospital, 60590 Frankfurt am Main, Germany; Institute of Biochemistry II, Goethe University, Frankfurt am Main, Germany; Department of Structural Biology, St. Jude Children’s Research Hospital, Memphis, USA; Department of Chemistry, Purdue University, West Lafayette, Indiana 47907, United States; Division of Experimental Neurosurgery, Department of Neurosurgery, University Hospital Heidelberg, INF400, 69120 Heidelberg, Germany; Buchmann Institute for Molecular Life Sciences, Goethe University Frankfurt, 60438 Frankfurt am Main, Germany; Institute for Clinical Pharmacology, Faculty of Medicine, Goethe University Frankfurt, Frankfurt am Main, Germany; Cardio-Pulmonary Institute, 60590 Frankfurt am Main, Germany; Department of Chemistry and Winship Cancer Institute, Emory University, Atlanta, Georgia 30022, United States; Frankfurt Cancer Institute, Goethe University, 60590 Frankfurt am Main, Germany; German Cancer Consortium (DKTK), Partner Site Frankfurt, 60590 Frankfurt am Main, Germany and German Cancer Research Center DKFZ, 69120 Heidelberg, Germany

**Author notes:** Radiation Biology and DNA repair, TU Darmstadt. (A.H.), (J.S.), (A.C.), (T.K.), (J. E.), (D. K.), (B. L.), (S. H.), (F. R.), (M.E.H.), (R.R.); (I.D.), (Ö.G.), (Y. W.), (X. Z.), (C.H.M.), (F.P.), (I.T.), (M.D.).

**Keywords:** Glioblastoma, Glioma Stem-Like Cells, BRAT1, Curcusone D, Invasion/Migration

## Abstract

Glioblastoma (GBM), the most malignant primary brain tumor in adults, has poor prognosis irrespective of therapeutic advances due to its radio-resistance and infiltrative growth into brain tissue. The present study assessed functions and putative druggability of protein breast cancer type 1 susceptibility protein (BRCA1)-associated Ataxia telangiectasia mutated (ATM)-activator 1 (BRAT1) as a crucial factor driving key aspects of GBM, including enhanced DNA damage response and tumor migration. By a stable depletion of BRAT1 in GBM and glioma stem-like (GSC) cell lines, we observed a delay in DNA double-strand break repair and increased sensitivity to radiation treatment, corroborated by *in vitro* and *in vivo* studies demonstrating impaired tumor growth and invasion. Proteomic analyses further emphasize the role of BRAT1’s cell migration and invasion capacity, with a notable proportion of downregulated proteins associated with these processes. In line with the genetic manipulation, we found that treatment with the BRAT1 inhibitor Curcusone D (CurD) significantly reduced GSC migration and invasion in an *ex vivo* slice culture model, particularly when combined with irradiation, resulting a synergistic inhibition of tumor growth and infiltration. Our results reveal that BRAT1 contributes to GBM growth and invasion and suggest that therapeutic inhibition of BRAT1 with CurD or similar compounds might constitute a novel approach for anti-GBM directed treatments.

## 1. Introduction

Tumors originating in the central nervous system pose significant challenges due to the delicate nature of the brain. The brain possesses unique physiological features such as the blood-brain barrier and specialized neuron and diverse glial cell types. Therefore, a phenotypically and genotypically distinction is made between diffuse or anaplastic astrocytomas (WHO grade 2 and 3), WHO grade 2 and 3 oligodendrogliomas, grade 4 glioblastomas (GBM) and diffuse pediatric gliomas. These entities are further divided into different subgroups according to their genotypic characteristics (Louis *et al*, 2016; Louis *et al*, 2021; Olar & Aldape, 2014; Verhaak *et al*, 2010). Among those cancers, GBMs represent the most aggressive malignant tumor type, with a median survival rate of only 15 months despite maximally possible treatment (Tran & Rosenthal, 2010; Becker & Yu, 2012). This is primarily caused by early relapses due to diffusely infiltrating tumor cells migrating as single cells or small clusters into the neuropil (network of intertwined neurons and glia cells). Therefore, patients cannot be completely cured by surgical resection of these tumors (Becker & Yu, 2012; Claes *et al*, 2007). GBMs are thus treated with a combination of radiation and chemotherapy (RCT) (Davis, 2016), despite being resistant to this multimodal treatment (Becker & Yu, 2012).

Treatment failures are mostly caused by the early filtration of tumor cells into the surrounding brain tissue, preventing complete tumor resection and leading to inevitable recurrence, largely attributed to treatment-resistant glioma stem-like cells (GSCs) (Bradshaw *et al*, 2016). The invasive behaviour of GBM cells is facilitated by complex interactions between tumor cells and the surrounding microenvironment. Signaling pathways such as the Mitogen-activated protein kinase (MAPK)/ extracellular signal-regulated kinase (ERK), phosphoinositide-3-kinase (PI3K)/ Akt/ mammalian target of rapamycin (mTOR; also known as the mechanistic target of rapamycin), and wingless/integrated (Wnt) pathways play critical roles in regulating cytoskeletal dynamics, cell adhesion, and extracellular matrix remodelling, thereby promoting GBM cell migration and invasion (Barzegar Behrooz *et al*, 2022). Additionally, transcription factors such as TWIST1 and SNAIL1 promote epithelial-mesenchymal transition (EMT), a process associated with increased invasiveness and stemness in GBM and other cancer entities (Myung *et al*, 2014; Fan *et al*, 2012). Furthermore, GBM cells have been observed to migrate along existing brain structures, including white matter tracts, blood vessels, and the extracellular matrix, facilitated by interactions with multiple transmembrane receptors. GSCs, distinguished by their increased differentiation potential compared to other tumor cells, typically reside within specialized niches and express various stemness-associated marker proteins. The abundance of GSCs, which possess enhanced migratory capacities and therapy resistance, further exacerbates the invasive behaviour of GBM (Mair *et al*, 2018; Bradshaw *et al*, 2016).

Further, it has been shown that GSCs are particularly resistant to radiotherapy due to mutations in the DNA damage response (DDR), which facilitate DDR and allow the tumor cells to survive in a quiescent state (Bao *et al*, 2006). According to the cancer stem cell model, most cancer cells can de-differentiate to a cancer stem-cell like phenotype, but can also become differentiated cancer cells again. Accordingly, one approach is to eliminate this type of cells from the tumor population to reduce the probability of a relapse (Dirkse *et al*, 2019). For this, we have recently shown that inhibiting the Hedgehog and NOTCH pathways in GSCs with Arsenic Trioxide (ATO) in combination with the anticancer agent (-)-Gossypol (Gos) which reduces not only stemness markers/properties, but also decreases expression of DNA repair and proteins involved in cell migration. One protein, which was prominently depleted in this study was the protein breast cancer type 1 susceptibility protein (BRCA1)-associated ATM activator 1 (BRAT1) (Linder *et al*, 2019). BRAT1 is ubiquitously expressed and interacts with BRCA1 and ATM and is therefore potentially involved in the DDR, cell proliferation, cell migration and invasion (Aglipay *et al*, 2006; So & Ouchi, 2013). Interestingly, BRAT1 variants are associated with diverse clinical conditions, from neonatal seizures to neurodevelopmental disorders (Fowkes *et al*, 2022).

The curcusone diterpenes, isolated from *Jatropha curcas*, share a tricyclic carbon skeleton with the daphnane and tigliane diterpenes and have shown promising anticancer properties, BRAT1 was identified as a key cellular target of the curcusones using chemoproteomics. Therefore, exhibiting antitumor effects are probably mediated at least in part by interfering with BRAT1/integrator subunit complexes (INTS11/INTS8) (Dokaneheifard *et al*, 2024) and thereby blocking promoter activity and likely transcription of BRAT1 itself. Curcusone D (CurD) leads to an impaired DDR, reduced cancer cell migration, and potentiated activity of the DNA damaging drug etoposide, in HeLa, MCF-7 and MDA-MB-231 cells, indicating an essential aspect in targeting cancer metastasis (Cui *et al*, 2021). This finding is significant as BRAT1 has been considered a non-pharmaceutical targetable oncoprotein with no known small molecule inhibitors.

Understanding the molecular mechanisms driving GBM invasion and migration is crucial for the development of novel therapeutic strategies. Moreover, advances in imaging techniques, like OTCxLSFM, provide valuable tools for studying GBM invasion dynamics and evaluating the efficacy of anti-invasive therapies (Haydo *et al*, 2023). By elucidating the complexities of GBM cell invasion and migration, may lead to the development of more effective therapeutic interventions for this devastating disease. Here, we have shown that BRAT1 plays a pivotal role in regulating cancer cell behaviour by impacting both migration and invasion, while also exerting influence over DDR. Moreover, we have identified BRAT1 as a novel critical factor contributing to the aggressive pro-migratory and pro-invasive characteristics of GBM. Lastly, we have identified the BRAT1 inhibitor CurD as a putative candidate that partially increases radio sensitivity of GBM and aims for the development of targeted therapy for GBM.

## 2. Results

### 2.1 BRAT1 expression is elevated in GBM

Recently, we applied a combination of two anticancer drugs, arsenic trioxide (ATO) and (-)-Gossypol (Gos, also known as AT-101), and showed that the combination leads to profound reduction of proteins involved in the DDR as well as 40% of the proteins which were downregulated cluster under the term movement. In this dataset, one of the most prominent protein which was downregulated was BRAT1 (Linder *et al*, 2019). An analysis of publicly available datasets revealed that BRAT1 is overexpressed in GBM compared to healthy tissues (Figure 1A) and it correlates negatively with patient survival according to the Gravendeel-dataset (Gravendeel *et al*, 2009) analyzed via the GlioVis portal (Bowman *et al*, 2017) (Figure 1B). Further, an analysis of the more recent CGGA database (Zhao *et al*, 2017) revealed that BRAT1 expression increases with tumor grade (Figure 1C). These data suggest that BRAT1 is an as-of-yet underappreciated, but important factor for GBM aggressiveness. To investigate the potential effects of BRAT1, stable BRAT1-knockdown (KD) cells were developed using lentiviral shRNA constructs for the GBM cell line U251 and the GSC line NCH644. The shBRAT1 cell lines showed a clear reduction in BRAT1-expression as compared to shCtrl cells (Figure 1D and 1E). Densitometric quantification further confirmed these findings. Residual BRAT1 expression in U251 cells, is approximately 10% for shBRAT1 (Figure 1D) and in NCH644 shBRAT1 cell line the remaining BRAT1-expression was around 25% (Figure 1E).

**Figure 1.**
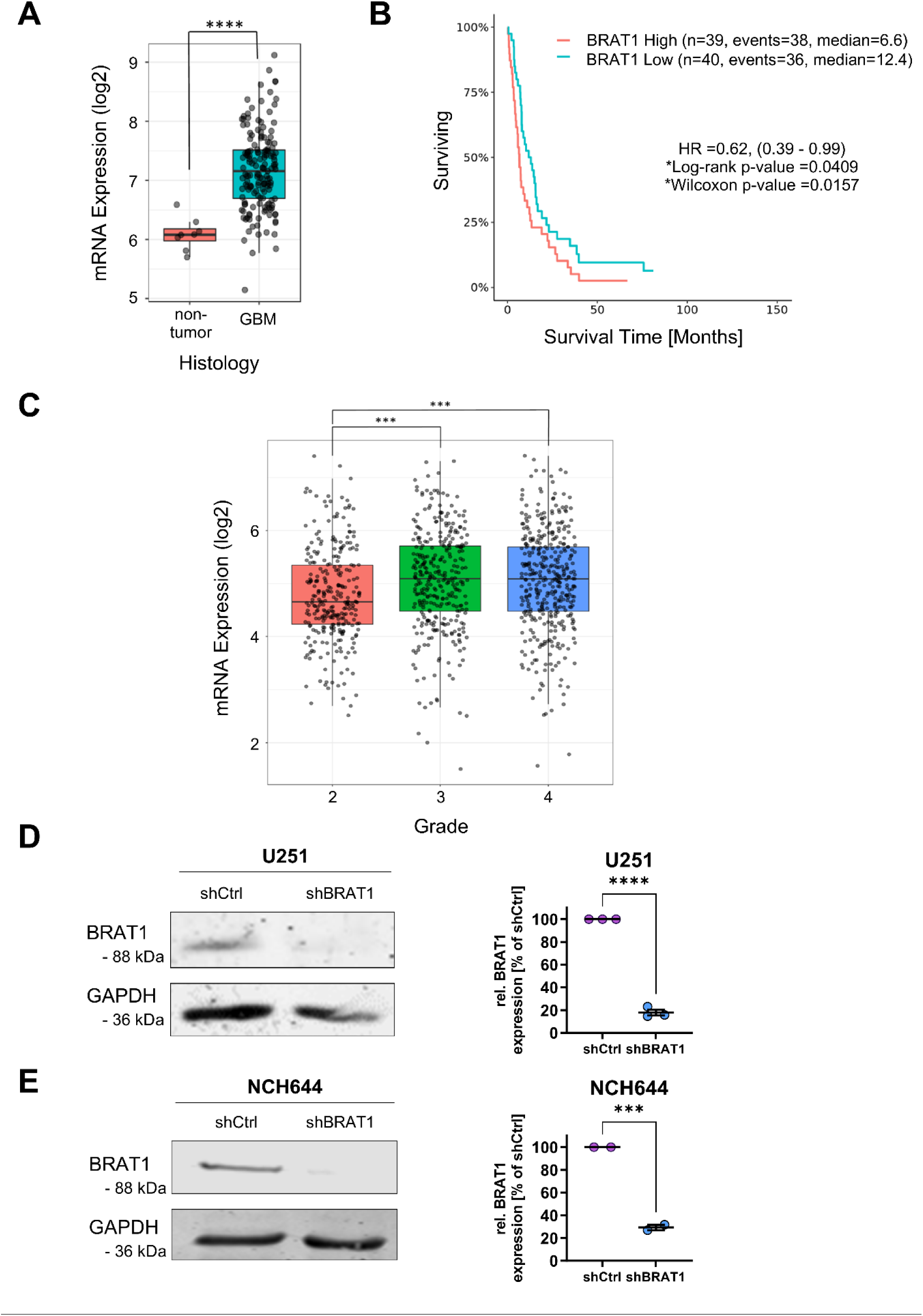
BRAT1 expression is increased in GBM. (A) Increased levels of BRAT1 compared to healthy tissue. Analysis of Gravendeel-dataset (Gravendeel et al, 2009) using the GlioVis portal (Bowman et al, 2017). (B) Higher BRAT1 levels are associated with decreased GBM patient survival. (C) According to the CGGA dataset (Zhao et al, 2017) BRAT1 expression is elevated in higher grade tumors. (D) Western blot and densitometric validation of stable shRNA BRAT1 depletion in U251 GBM cells, GAPDH as housekeeping protein; Scatters show biological replicates. (E) Western blot and densitometric validation of stable shRNA BRAT1 depletion in NCH644 GSCs; GAPDH as housekeeping protein; Scatters show biological replicates. Statistics: A: Unpaired two-sided t-test between group levels; B: Survival analysis using Wilcoxon signed rank test. C: One-way ANOVA with posthoc t-test using a correction of alpha according to Dunnett versus control. D, E: Unpaired two-sided t-test between groups. Error bars are SEM. *: p < 0.05; ***: p < 0.001; ****: p < 0.0001.

### 2.2 BRAT1 is essential for an efficient DNA repair

Since BRAT1 expression was previously shown to be increased in radioresistant GBM cells and GSCs (Linder *et al*, 2019), we next examined the DNA repair proficiency in BRAT1 KD cells using γH2AX and 53BP1 foci assays. For this, the cells were irradiated with 10 Gy (U251 GBM cells) and 8 Gy (NCH644 GSCs). At 1 hour (h) and 24 h after irradiation (IR), the cells were fixed and immunostained for γH2AX and 53BP1 to visualize and quantify DNA double strand breaks (DSBs) (Figure 2). The same approach was performed in BRAT1 overexpressing cells (Suppl. Figure 1).

**Figure 2.**
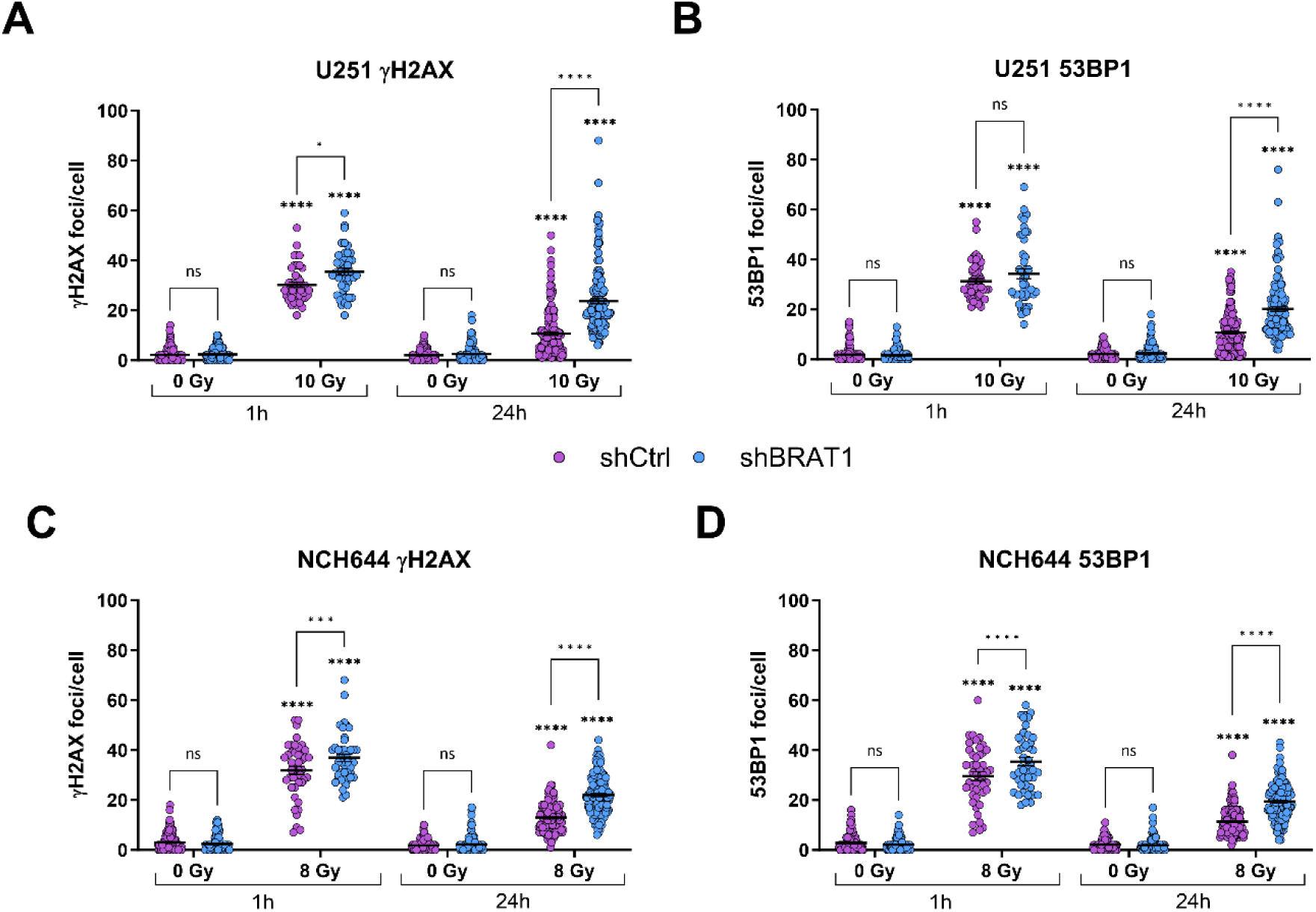
BRAT1 is essential for efficient DNA repair. shCtrl and shBRAT1 cells were compared in two cell lines. (A) U251 γH2AX foci assays and (B) U251 53BP1 foci assays 1 h and 24 h after 10 Gy IR. (C) NCH644 γH2AX foci assays and (D) NCH644 53BP1 foci assays 1 h and 24 h after 8 Gy IR. At 1 h after IR γH2AX- or 53BP1-positive foci were counted in 15 cells per condition; in non-IR cells and 24 h after IR, foci in 50 cells were counted per condition. The experiment was performed 2 times in biological replicates and results were pooled with overall n≥15. Statistics: Two-way ANOVA with Tukey’s multiple comparisons test. Error bars are SEM. ns = not significant; * p < 0.05; *** p < 0.001; **** p < 0.0001 against respective non-IR cells or as indicated.

After, 1 h of 10 Gy IR, both U251 shCtrl and U251 shBRAT1 cells showed similar numbers of DNA DSBs of around 30-40 foci/cell (Figure 2A). However, at 24 h after IR, the U251 shCtrl cells displayed approximately 10 foci/cell, whereas the U251 shBRAT1 cells indicated significantly higher DNA DSB numbers of around 25 γH2AX foci/cell. These findings were confirmed by the 53BP1 staining with 10 foci (shCtrl) and 22 foci (shBRAT1), respectively (Figure 2B). To further validate our observation, we tested the same setup on the GSC line NCH644 (γH2AX: Figure 2C and 53BP1: Figure 2D) resulting comparable increased residual foci numbers upon BRAT1 knockdown. To further test our hypothesis that BRAT1 is needed for efficient DNA repair, we implemented the same setup of γH2AX and 53BP1 staining in U251 and NCH644 cells transduced with V-lentiCMV-Puro (empty vector control) and V-lentiCMV-hBRAT1-Puro for establishing BRAT1-overexpressing (BRAT1-OE) cells. BRAT1 overexpression was validated using western blot analyses (Suppl. Figure 1A). Indeed, we could observe reversed effects in comparison to BRAT1-deficient cells, showing decreased γH2AX and 53B1 foci/cell in GBM U251 (Suppl. Figure 1B) and GSC NCH644 BRAT1-OE cells (Suppl. Figure 1C), indicating a more efficient DDR.

### 2.3 Knockdown of BRAT1 decreases migration *in vitro* and *in vivo*

To analyze the impact of BRAT1 expression on the migration potential in a cell-based model, we performed a wound healing assay to analyze the gap closure of U251 GBM cells and performed a transwell migration assay for NCH644 GSC. In addition, we used an orthotopic GBM mouse model to confirm the essential role of BRAT1 in promoting tumor survival (Figure 3). First to assess *in vitro* migration U251 shCtrl and shBRAT1 cells were seeded in IBIDI inserts, creating a defined 500 µm gap in between two adherent colonies. This allowed the monitoring of the migration capacities to finally observe the gap closure (%). Overall, the readout was the percentage of gap closure up to 72 h of culture (Figure 3A) indicating a deceased migration potential of U251 shBRAT1 cells of about 60% compared to U251 shCtrl with approximately 80% gap closure (Figure 3B). Similar findings were observed in NCH644 BRAT1-depleted cells fixed after 48 h incubation (Figure 3C), with around 400 migrated cells compared to shCtrl, with approximately 440 migrated cells (Figure 3D). To further asses the *in vivo* relevance and translation, we implanted NCH644 shBRAT1, as well as BRAT1-expressing NCH644 shCtrl into athymic nude mice and observed clinical development (scores and body weights) and mouse survival over 12 weeks. BRAT1-depletion significantly enhanced median overall survival from 42.5 days of shCtrl NCH644 tumor-bearing mice to 55 days indicating that BRAT1 depletion prolonged survival (Figure 3D). To further test our results, we performed another migration assay, using U251 empty vector control cells and U251 BRAT1-OE cells (Sup. Figure 2). A reversed effect was observed. By this, BRAT1-OE cells depicted an overall higher gap closure over the course of 72 h (Suppl. Figure 2A). This was further confirmed by the finding that BRAT1 overexpression led to a 90 to 100% gap closure, compared to empty vector controls with 60 to 80% gap closure after 72 h (Suppl. Figure 2B).

**Figure 3.**
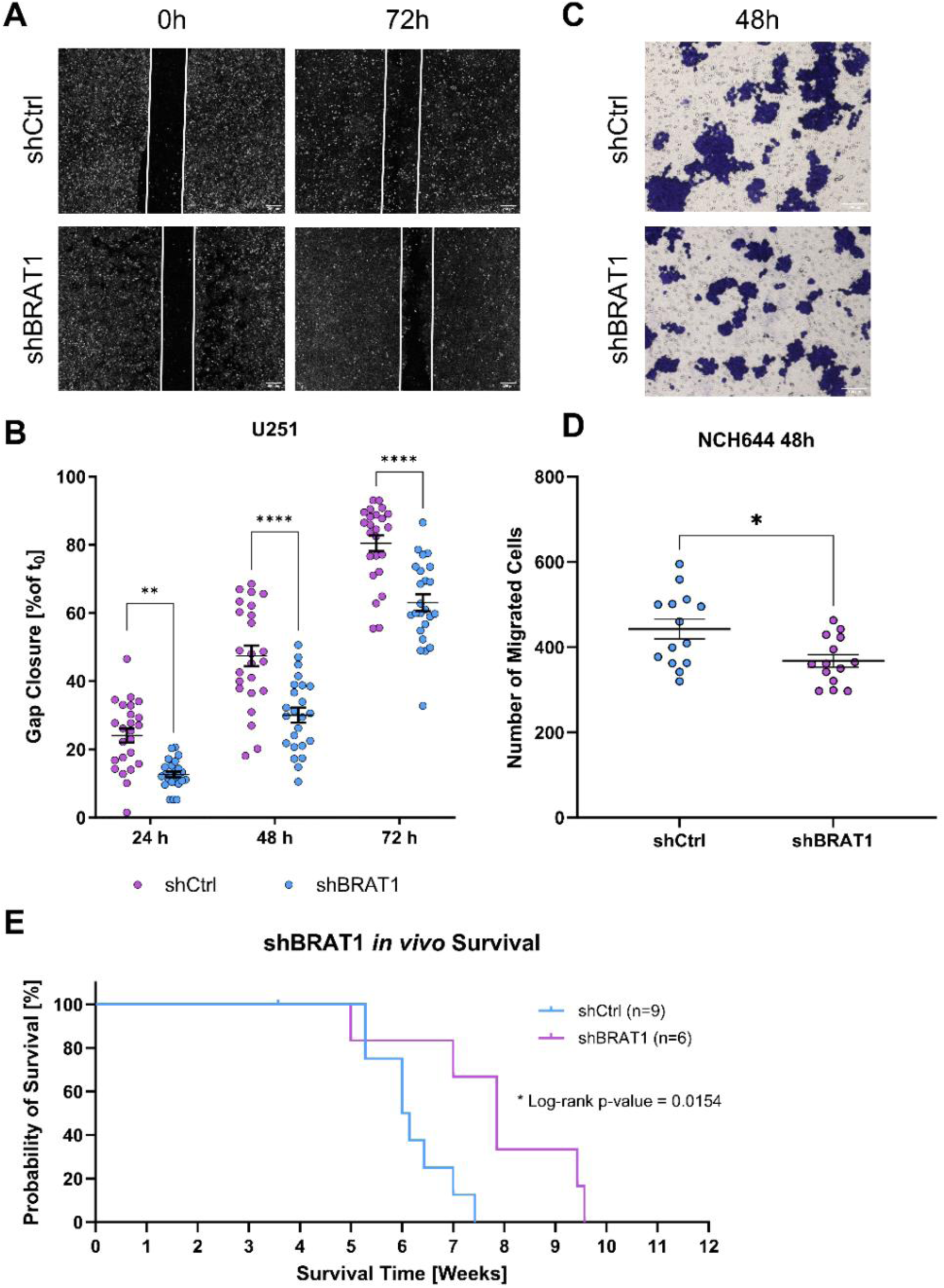
BRAT1 knockdown reduces migration in vitro and prolongs survival of NCH644 GSCs in an orthotopic transplantation model. (A) IBIDI migration assay pictures depicting U251 shCtrl and U251 shBRAT1 gap closure after 72 h; scale bar 200 µm (B) Quantification of U251 shCtrl and U251 shBRAT1 IBIDI migration assay displaying 24 h, 48 h and 72 h of gap closure in %. The experiment was performed 3 times in biological replicates, results were pooled with an overall n = 8 (C) Transwell migration assay images depict NCH644 shCtrl and NCH644 shBRAT1 fixed after 48 h; scale bar = 200 µm. (D) Quantification (number of migrated cells) in transwell migration assay at 48 h. The experiment was performed 3 times in biological replicates, results were pooled with an overall n≥4. (E) Kaplan-Meyer survival curve of tumor bearing mice. Mice were orthotopically implanted with either NCH644 shCtrl (n = 9 mice) or NCH644 shBRAT1 (n = 6 mice), observed over the course of 12 weeks. Statistics: B, D: Unpaired two-tailed t-test with Mann–Whitney test. E: Survival was assessed with Wilcoxon signed log rank test. Error bars are SEM, * p < 0.05; ** p < 0.01; **** p < 0.0001 against respective shCtrl cells as indicated.

In addition, we assessed if BRAT1 influenced the stemness potential of GSCs, and therefore analysed the ability to form spheres in suspension cultures (Roth *et al*, 2024). For this purpose, we used NCH644 GSCs. In that context, shCtrl versus shBRAT1 and empty vector control cells versus BRAT1-OE cells. After incubation for 7 days (d) the cells were analyzed with a Tecan Spark plate reader and sphere number (Suppl. Figure 3A) and sphere area (Suppl. Figure 3B) was determined using Fiji ImageJ software (Schindelin *et al*, 2012). As given in supplemental figure 3, BRAT1 depletion leads to less sphere numbers (∼20 spheres) compared to shCtrl cells (∼70 spheres), whereas BRAT1 overexpression (∼100 spheres), leads to an increased sphere number compared to empty vector control cell line (∼70 spheres). However, the sphere area is only reduced in BRAT1-depleted cells (∼ 600 µm) compared to shCtrl (∼800 µm), whereas BRAT1 overexpression (∼710 µm) does not influence the sphere area compared to its empty vector control (∼700µm) (Suppl. Figure 3B).

### 2.4 Global proteomic- and phospho-proteomic data reveals that proteins related to migration and invasion are downregulated upon BRAT1 knockdown

In order to gain insights into the role of BRAT1 in GBM cells and GSCs we next performed a proteomic and phospho-proteomic analysis of BRAT1-depleted cells. Firstly, the global total proteome of BRAT1-depleted cells indicates changes in proteins associated with cancer cell migration and invasion in line with the findings from our previous experiments. Our analysis of U251 BRAT1-depleted cells, revealed a total of 7643 proteins (Figure 4A) with 77 proteins to be significantly increased and 96 to be significantly downregulated (including BRAT1) in comparison to shCtrl cells using distinct cut offs p≤0.05 and log-fold changes greater than 0.5. Among the most strongly decreased, we observed proteins related to DNA repair (FANCA, USP38, NET1), autophagy (BCAS3), cancer cell migration and/or invasion (RASSF2, KMO, BCAS3, GNA12), as well as proteins that can be generally considered as pro-tumorigenic, particularly in glioma (KMO, BCAS3, GNA12) (Figure 4A). Exemplarily, Ras association domain family member 2 (RASSF2), which is a member of the Ras association domain family implicated in various signaling pathways including the Hippo signaling pathway or Ras signaling pathway, which regulate cell migration and invasion is downregulated by a factor of 2.3 (Dhanaraman *et al*, 2020). On the other hand, among the increased proteins are proteins involved in the blockade of cancer cell migration and invasion (SCAI, CARMIL2), autophagy activation (HSBP1, AMBRA1, NBR1, RILP) as well as proteins that can be considered as anti-tumorigenic (SCG2, DIP2A, HSBP1) (Figure 4A). Overall, these data suggest that depletion of BRAT1 evokes changes that go beyond the DDR and includes migratory and invasive capacity changes as well as metabolic changes like fatty acid oxidation.

**Figure 4.**
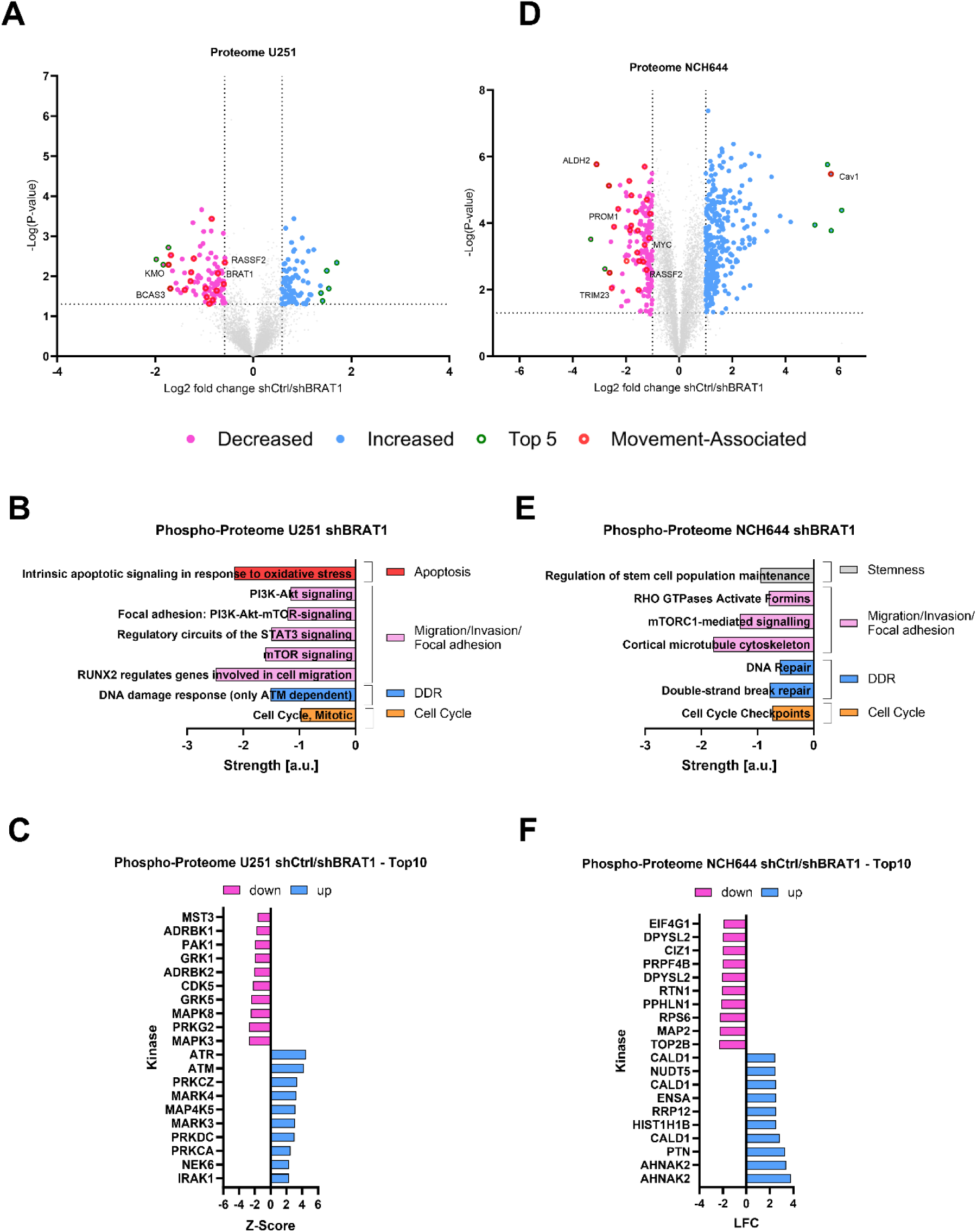
Proteomic and phospho-proteomic analysis suggest that BRAT1 influences migration and invasion in U251 GBM cells and NCH644 GSCs. (A) Volcano Plots of proteomic data show significantly decreased (pink) or increased (blue) and the top 5 most strongly (green dots) changed proteins upon BRAT1 knockdown. Proteins related to “movement” in shCtrl/shBRAT1 U251 GBM cells are depicted in red dots. (B) Pathway enrichment of decreased proteins using STRING software in U251 cells after BRAT1-depletion, depicted in overall strength. Decreased pathways were associated with the terms apoptosis, cell migration/invasion/focal adhesion, DDR and cell cycle. (C) Kinase-Substrate-Enrichment-Analysis (KSEA) of U251 shCtrl and shBRAT1 cells suggests that BRAT1-depletion hinders migration and advancing through proceeding of the DNA damage response. Phospho-Proteomic analysis of the Top10 suggests a loss of migratory potential and differential regulation of MAPK-pathway in BRAT1-depleted cells. (D) Volcano Plots of proteomic data show significantly decreased (pink) or increased (blue) and the top 5 most strongly (green dots) changed proteins upon BRAT1 knockdown. Proteins related to “movement” in shCtrl/shBRAT1 NCH644 GSCs are depicted in red dots. (B) Pathway enrichment of decreased proteins using STRING software in NCH644 cells after BRAT1-depletion, depicted in overall strength. Decreased pathways were associated stemness maintenance, cell migration/invasion/focal adhesion, DDR and cell cycle. (F) Top10 Kinases of NCH644 shCtrl and shBRAT1 cells indicate that BRAT1-depletion hinders migration and prevents proceeding of the DNA damage response, loss of migratory potential as well as influence on Rac/Rho- and MAPK-pathways.

Using STRING software (Szklarczyk *et al*, 2019) a distinct selection of decreased pathways after BRAT1-depletion in U251 GBM cells was performed (Figure 4B). Using this, BRAT1 can be possibly associated by negatively influencing pathways associated with migration/invasion and focal adhesion like: mTOR signaling, signal transducer and activator of transcription 3 (STAT3) signaling pathways and focal adhesion using PI3K-Akt-mTOR-signaling pathway. Besides this, BRAT1 may influence pathways associated to apoptosis and cell cycle.

To confirm our findings, we analysed the Top10 decreased (pink) and enriched (blue) kinases upon BRAT1 depletion (Figure 4C). Further, using the Perseus software for the phosphoproteomic data set of U251 GBM cells after BRAT1 depletion, a kinase-substrate enrichment analyses (KSEA) of phosphoproteomic data sets after BRAT1 depletion was performed. The phospho-proteomic content of BRAT1-depleted cells pointed towards a role for sustained DNA damage response, autophagy and migration. Kinase-substrate predictions were inferred from phosphoproteomic data sets of U251 GBM cells after BRAT1 depletion using the KSEA-App (https://casecpb.shinyapps.io/ksea/). Kinase-substrate searches included PhosphoSitePlus and NetworKIN with scores 2 or greater. After bioinformatic analyses of phospho-proteomic data of U251 cells upon BRAT1-depletion, the two most active kinases were ATM and ATR, as well as DNA-dependent protein kinases (DNA-PKcs), which would suggest a baseline activation of the DDR. However, the most decreased kinases with z-scores -2, were MAPK3, also known as ERK1 and MAPK8, also known as JNK1, which are both members of the MAPK signaling pathway, which regulates various cellular processes including cell migration and invasion (Huang *et al*, 2004). Moreover, Cyclin-Dependent Kinase 5 (CDK5), involved in cytoskeletal dynamics, cancer cell migration and invasion and p21-Activated Kinase 1 (PAK1) that regulates cytoskeletal reorganization and cell motility, were among the phosphoproteins being downregulated (Martínez-Montemayor *et al*, 2010; Sharma *et al*, 2002; Bisht *et al*, 2015; Zhou *et al*, 2014) (Figure 4C). Lastly, proteins regulating MAP2K/MAPK-signaling, senescence and cell cycle-associated processes, were decreased, suggesting a loss of a pro-migratory phenotype (Bisht *et al*, 2015; Zhou *et al*, 2014).

The same analysis pipeline was applied for the GSC line NCH644 upon BRAT1-depletion, besides only displaying the overall decreased kinases without KSEA incorporation (Figure 4D-F). The total global proteome (7273 proteins) suggested that 10% of proteins which were downregulated (total 179 decreased proteins) are involved in movement, therefore influencing cancer cell migration and invasion. The top downregulated protein was aldehyde dehydrogenase 1 (ALDH1) which has been implicated in cancer stem cell maintenance and drug resistance and is associated with tumor invasion and metastasis (Wang *et al*, 2021). A similar function displays another protein Prominin 1 (PROM1), which was also amongst the most decreased proteins (Adamski *et al*, 2017). Another decreased protein was the tripartite motif containing 23 (TRIM23), which contributes to carcinogenesis in colorectal cancer and act as a poor prognostic factor (Han *et al*, 2020). Finally, RASSF2 was amongst the most decreased proteins, implying the importance of BRAT1 potentially influencing cancer cell invasion. Interestingly and further supporting our findings from the U251 proteome analysis, Caveolin-1 (Cav1) was one of the most upregulated proteins. Cav-1 has been reported to have complex effects on cancer cell migration and invasion, with both pro- and anti-migratory functions reported in different cancer types (Núñez-Wehinger *et al*, 2014) (Figure 4D).

To continue, using STRING software (Szklarczyk *et al*, 2019), a distinct selection of decreased pathways after BRAT1-depletion for NCH644 GSCs was performed (Figure 4E). Importantly, we could detect decreased pathway associated to stemness maintenance. This supports the findings from our stemness assay, in which BRAT1 knockdown/overexpression influences stemness capacity. Moreover, mTORC1-mediated signaling and RHO GTPases Activate Formins signaling pathways were decreased after BRAT1 depletion, supporting the role of BRAT1 negatively regulating migration and invasion and further validating the findings of the proteome data similar to the chemoproteomics of (Cui *et al*, 2021). Similar to the findings of U251 pathways like DDR, including efficient DNA double-strand repair and cell cycle were as well affected in NCH644 after BRAT1 depletion.

Lastly, to analyze the phospho-proteomic changes of NCH644 upon BRAT1-depletion a Top10 of most significantly down- and upregulated kinases were selected (Figure 4F). Interestingly, one of the most top upregulated kinases was Neuroblast differentiation-associated protein (AHNAK2), which is a large scaffolding protein that is primarily found in the cytoplasm and nucleus of cells. On the other hand, the top decreased kinases indicate downregulation of MAPK- and Rac/Rho-pathways upon KD of BRAT1. Topoisomerase II Beta (TOP2B) was the most significant downregulated kinase with a LFC greater than -2. TOP2 family members are involved in cell migration, invasion and EMT in cervical cancer via activating the PI3K/AKT signaling (Uusküla-Reimand & Wilson, 2022). Dihydropyrimidinase-like 2 (DPYSL2, also known as CRMP2) was described to be involved in cytoskeletal dynamics and has been associated with downstream effectors like ERK1/2 (Pham *et al*, 2016; Conde & Cáceres, 2009). Lastly one of the top downregulated kinase is Eukaryotic Translation Initiation Factor 4 Gamma 1 (EIF4G1), which can be involved in translation initiation and has been linked to cancer progression, including migration and invasion (Martínez-Montemayor *et al*, 2010; Tan *et al*, 2019; Jaiswal *et al*, 2019). All mentioned proteins/kinases and their functions are again listed for better oversight in supplemental figure 4.

### 2.5 Curcusone D, a BRAT1 inhibitor, decreases the pro-migratory/invasive phenotype of GBM and synergizes with irradiation

A study in 2021 isolated and characterized new curcusone diterpenes, which have shown promising anticancer properties and interestingly BRAT1 was identified as a key cellular target of the curcusones using chemoproteomics, however mechanistically unresolved and so far not reproduced by others. Curcusone D likely has off-target effects, but nonetheless served as a pharmacological tool in the present study to question and confirm the results obtained with BRAT1 knockdown cells (Cui *et al*, 2021). First, migration *in vitro* assays were performed on a variety of GBM/GSCs lines, analyzing the growth-inhibitory effects evoked by BRAT1 inhibition (Suppl. Figure 5). By using wound healing assays, we could confirm, that 3 µM CurD time-dependently blocks migration of 2D cell cultures (Suppl. Figure 5A), similarly as genetic BRAT1-depletion. Likewise, migration of the GSC line GS-5 (Günther *et al*, 2008) on laminin-coated plates was inhibited by CurD using a sphere migration assay (Haydo *et al*, 2023) (Suppl. Figure 5B). Further, migration of NCH644 GSCs using transwell migration assays was reduced under CurD-treatment (Suppl. Figure 5C). Taken together, these data demonstrate that BRAT1 inhibition via CurD effectively blocks migration *in vitro*. We next validated that CurD leads to enhanced numbers of DSBs and measured γH2AX fluorescence intensity via flow cytometry in both U251 GBM cells and NCH644 GSCs (Suppl. Figure 5D). In U251 GBM cells (Suppl. Figure 5D, left panel), 24 h treatment with CurD mildly increased the amount of γH2AX even in the absence of radiation. Shortly after a 10 Gy irradiation (1 h) the fluorescence intensity was significantly increased and further enhanced under CurD-treatment. 24 h post-10 Gy IR the γH2AX fluorescence intensity had recovered to baseline values in DMSO-treated U251 GBM cells, whereas treatment with CurD still had significantly increased DSB fluorescence signal. Similar results were obtained in NCH644 GSCs (Suppl. Figure 5D, right panel), although at 24 h post-8 Gy IR the remaining amount of γH2AX fluorescence intensity was only slightly higher compared to DMSO-treated GSCs.

Based on the above-presented data we inferred that 1) BRAT1-depletion might hinder tumor growth in more complex model systems and 2) BRAT1-depletion might also negatively impact tumor cell migration/invasion. In order to address both question we used an *ex vivo* approach consisting of organotypic tissue slice cultures (OTCs) of adult murine brains unto which GFP-positive tumor spheres are transplanted (Haydo *et al*, 2023; Gerstmeier *et al*, 2021; Linder *et al*, 2019; Remy *et al*, 2022). Fluorescently labeled tumor cells, in this case NCH644^GFP^ GSCs (Haydo *et al*, 2023) were placed onto murine brain slices and tumor development up to 17 d was monitored. We applied 5 µM CurD to the established tumors on the OTCs at d0, which was followed by a fractionated radiation protocol consisting of 3 times of 2 Gy for two weeks. In detail, every Monday, Wednesday and Friday cells were treated and irradiated, leading to a total 6 times 2 Gy IR dose over the course of two weeks. The culture medium containing CurD was renewed on the day of radiation 2-3 h prior to IR. Exemplary images are displayed in Figure 5A. The treatment with CurD caused a change in morphology of the GSCs compared to the control and the combination of CurD and IR resulted in an almost completely eradication of the tumor. Further, quantification of the tumor area confirmed the morphologic assessment (Figure 5B). Solvent control (DMSO-treated) and non-irradiated tumors grew continuously over the observation period and reached more a 30-fold increased size compared to d0. CurD-treatment and IR alone evoked a similar growth-suppressing effects. Lastly, the combination of CurD and IR led almost to a complete elimination of the tumors. To assess more information of this model, tumor area of specific timepoints were further quantified (Figure 5C). Specifically, in CurD treated tumors, the tumor increased in size until approximately day 7 and then reached a plateau. Hence, IR-treated tumors were significantly smaller at day 7 and reached a plateau at day 10. These results suggest that a combination would possibly potentiate the effects of both treatments. As such the tumors increased in size until day 5 and declined in size afterwards. Strikingly at the end of the experiment (d17) 30% of the tumors (8 of 24) were completely eliminated (Figure 5C). CurD-treatment led to tumors with distinctly clear-cut borders, suggesting complete failure to invade the brain tissue. Irradiated tumors appeared similarly invasive to solvent-treated tumors, but with decreased sizes. To confirm this observation we applied an imaging technique combining OTC culture with light sheet fluorescence microscopy (LSFM) called OTCxLSFM (Haydo *et al*, 2023) to measure the tumors and its particular single cell invasion in a 3D manner and display it in 90° angles (Figure 5D). This advanced microscopy approach confirmed that DMSO-treated tumors display diffuse 3D infiltration and spreading of cells into the surrounding brain tissue. In contrast, CurD-treatment devoided tumor migration. The quantitative assessment using the invasion distance measured from the tumor center as a readout confirmed the differences of tumor invasion, by decreased migration distance in CurD treated tumors with a maximal migrated distance of 1450 µm compared to solvent control cells with a maximal migrated distance of up to 1750 µm (Figure 5E, left panel). Whereas, the frequency distribution of such distances revealed the overall diffusion of the cells. Both groups displayed an average migrated distance of approximately 500 µm in solvent control cells and CurD treated tumors. However, in solvent control cells about 15% met this distance whereas only about 6% of CurD treated reached this distance (Figure 5E, right panel). Our three-dimensional approach revealed a reduction in tumor cell spreading and frequency distribution, again indicating a decreased migration potential and invasiveness by blocking BRAT1 using CurD.

**Figure 5.**
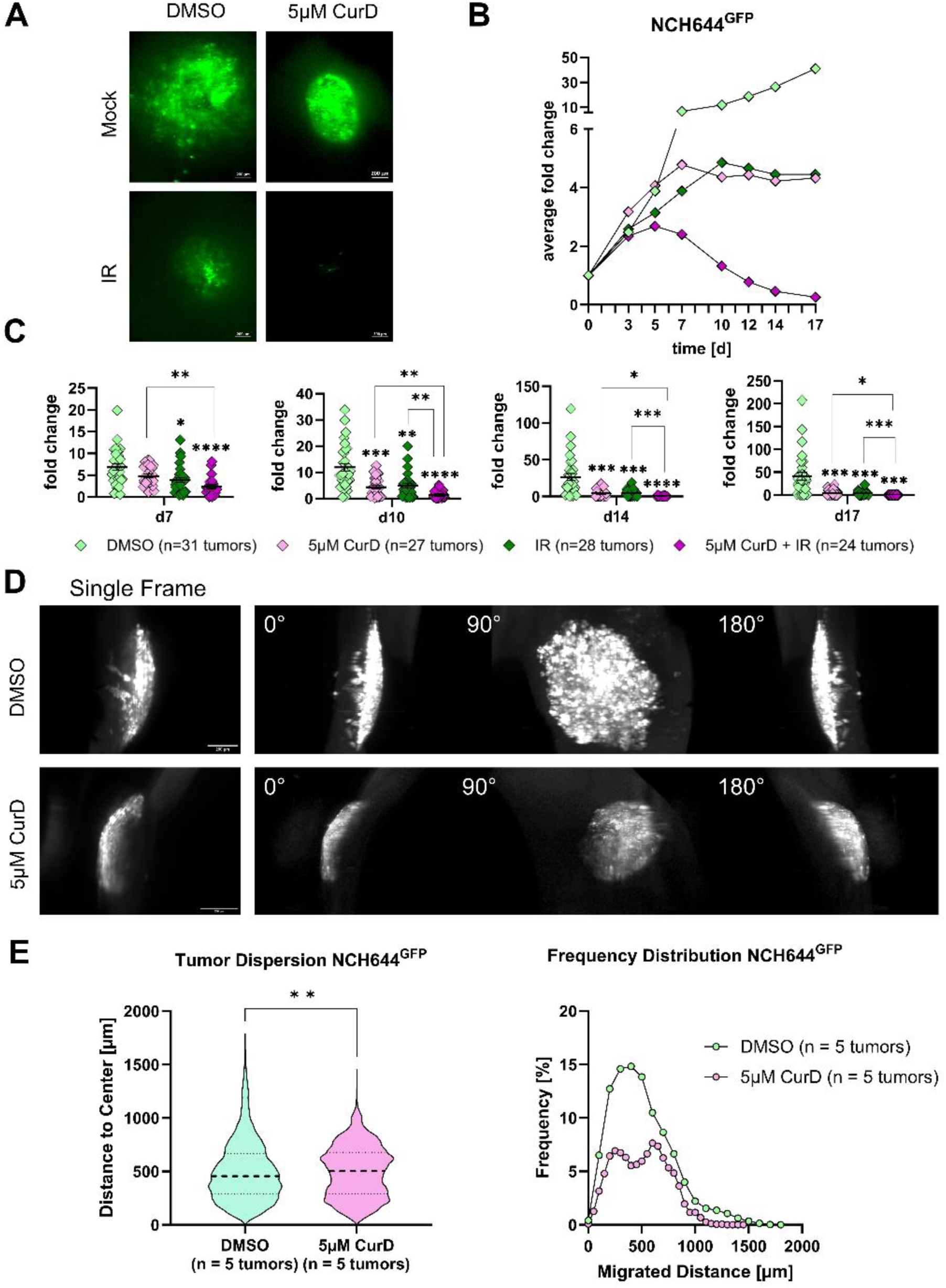
BRAT1 inhibitor Curcusone D inhibits tissue invasion and migration in an ex vivo OTC model and synergizes with IR-treatment. NCH644^GFP^ were used in this experimental setup and were treated with 5 µM CurD monotherapy and in combination with fractionated radiation therapy with 3 fractions of 2 Gy for two weeks. The culture medium containing CurD treatment was renewed on the day of radiation 2-3 h prior IR. (A) Representative stereomicroscopic tumor images of GSC NCH644^GFP^ after treatment on day 17. Scale bars = 200 µm. (B) Averaged growth kinetic throughout the observation time (timeline) of NCH644^GFP^ cells treated with DMSO as solvent control, 5 µM CurD, IR and combined treatment (5 µM CurD + IR). (C) Scatter plots show tumor growth at specific timepoints (d7, d10, d14 and d17) of the experiment shown in B. (D) The montage of 3D reconstruction LSFM images show the same tumor at the latest time point (d17) of incubation for NCH644^GFP^ treated with DMSO or 5 µM CurD displayed in a single frame and 90° angles. Scale bars = 200 μm. (E) Evaluation of the LSFM pictures for NCH644^GFP^ (DMSO n = 5 tumors and 5 µM; CurD n = 5 tumors; being biological replicates) at the latest time point (d17). Violin plot depicting tumor dispersion and frequency distribution showing the relative amount of migration distance in 500 μm bins of DMSO n = 5 tumors and 5 µM; CurD n = 5 tumors; being biological replicates at the latest time point (d17). Statistics: C: Two-way ANOVA with Tukey’s multiple comparisons test. E: Unpaired t-test. Error bars are SEM. * p < 0.05; ** p < 0.01; *** p < 0.001; **** p < 0.0001 against respective DMSO treated cells or as indicated.

## 3 Discussion

GBM, the most malignant primary brain tumor in adults, is usually treated with combined RCT following surgery according to current standards, despite its pronounced radio- and chemoresistance (Becker & Yu, 2012). GBM further exhibits a strong pro-migratory and pro-invasive features (Hanahan & Weinberg, 2011; Hatoum *et al*, 2019). It is believed that the persistence of GSCs in this tumor is one of the major causes for treatment resistance, intra-tumoral heterogeneity and the pro-migratory/invasive phenotype. These GSC cells can self-renew infinitely, upregulate DNA damage repair pathways and escape radio- and chemotherapy by surviving in a quiescent state (Bao *et al*, 2006; Chen *et al*, 2012). In addition, these cells show a high capacity of cell invasion and cell migration (Haydo *et al*, 2023). Our previous proteomic approach revealed that BRAT1 is one of the most prominently decreased proteins after combined in vitro treatment of GSCs with ATO and Gos. Similarly this treatment inhibited tumor growth in our OTC *ex vivo* model, accompanied by reduced levels of proteins clustered under the term movement (40% of total decreased proteins), as well as a decrease in the levels of key DDR components (Linder *et al*, 2019). These findings raised the questions whether BRAT1 might play a functional role in GBM’s resistance to DNA damage and regulation of GBM migration/invasion. Of note, BRAT1 mutations are linked with neurodegenerative diseases and neurodevelopmental disorders such as rigidity and multifocal-seizure syndrome, although the exact molecular mechanisms underlying disease pathology are not well understood (Kong *et al*, 2024). Recently, it was shown that INTS11 and INTS9 subunits of the Integrator complex interact with BRAT1 and form a trimeric complex in human HEK293T cells as well as in the pluripotent human embryonal carcinoma cell line (NT2) to regulate the expression of neurodevelopmental genes (Dokaneheifard *et al*, 2023). A further study showed that the interaction of BRAT1 and the subunits of the Integrator complex (INTS9/INTS11) are responsible for processing the 3’ ends of various noncoding RNAs and pre-mRNAs (Cihlarova *et al*, 2022).

Importantly, CurD was described to reduce the expression of BRAT1 through unclear mechanisms but providing a pharmacologic tool to further assess the role of BRAT1 in GBM growth (Cui *et al*, 2021). Expression of BRAT1 is higher in grade 4 gliomas compared to other high-grade gliomas and positively correlated with decreased overall patient survival not only in glioma, but also in renal and liver cancer, identifying BRAT1 as a putative new target for cancer treatment (https://www.proteinatlas.org/ ENSG00000106009-BRAT1/pathology). To study the functional role of BRAT1 in GBM, shRNA-mediated BRAT1 depletion was used as a strategy to investigate various aspects of cancer pathophysiology and the cellular radiation response. To address the potential relevance of BRAT1 for the DDR, the time-dependent efficiency of radiation-induced DNA repair was analysed, as BRAT1 has been described to interact with ATM and BRCA1, two fundamental proteins in the early DDR (Jean Af & Toru Ouchi, 2016). Based on the assumption that BRAT1 1) is needed for ATM phosphorylation, 2) is needed for the function of BRCA1 and 3) interacts with DNA-PKcs, we hypothesized that DNA DSBs introduced by IR may not be repaired adequately in cells depleted of BRAT1, while overexpression of BRAT1 will result a reverse effect on the DDR (Low *et al*, 2015; So & Ouchi, 2011; Aglipay *et al*, 2006). In U251 GBM cells, shCtrl and shBRAT1 cells showed equal levels of DSBs 1 h after IR with 10 Gy. However, 24 h after IR, almost all IR-induced DNA DSBs were repaired in the control cells, but significantly elevated levels of DSBs were evident in BRAT1-depleted cells, indicating a defective DNA repair. This was also true in NCH644 GSCs, where we additionally observed that already 1 h after IR with 8 Gy significantly more DSBs were not fully repaired in the cells with depleted BRAT1. Further, BRAT1 KD has been shown to decrease phosphorylation of ATM and CHK2, crucial for effective DNA repair, as well preventing the premature dephosphorylation and detachment of ATM and DNA-PKcs from DNA damage sites (So & Ouchi, 2011; So & Ouchi, 2013). Experiments revealed that BRAT1-depleted cells have more pATM- and 53BP1 positive foci due to impaired DNA repair. Further, TOP2B generating transient DNA double-strand breaks to relieve topological stress during DNA replication and transcription was the most decreased kinase after BRAT1 depletion supporting the interaction of BRAT1 and DDR associated kinases (Broderick *et al*, 2019; Papapietro & Nejentsev, 2022). Moreover, treatment with the BRAT1 inhibitor CurD increased the foci, suggesting that BRAT1 was essential for sustained DDR activation. Additionally, here after IR, pATM-positive foci were higher in BRAT1-KD and CurD-treated cells compared to controls, highlighting BRAT1’s role in maintaining long-lasting DDR. In parallel, (Low *et al*, 2015) could show that BRAT1 is required for increasing the abundance of pATM by Nedd4 Family Interacting Protein 1 (Ndfip1). The results also suggested that BRAT1 was more crucial for sustained ATM phosphorylation in GSCs than in adherent GBM cells, indicating a potential therapeutic target for enhancing DNA repair efficacy.

To further investigate the pro-migratory phenotype of GBM and GSC lines (Seker-Polat *et al*, 2022), we tested *in vitro* migration to obtain insights into the migration capacity in relation to BRAT1 proficiency vs deficiency. Here we could demonstrate that genetic BRAT1 depletion leads to a decreased migratory potential compared to shCtrl cells. In an orthotropic GBM mouse model we could prove that BRAT1 depletion prolonged overall survival, in line with evidence suggesting that BRAT1 is an unfavourable prognostic marker in several cancers (https://www.proteinatlas.org/ENSG00000106009-BRAT1/pathology). However, there was no overall difference in the morphology of tumor sections as assessed by histochemistry. The pro-migratory role of BRAT1 was further validated by global proteomic and phospho-proteomic approaches, where proteins associated not only with migration (RASSF2, KMO), but also proteins associated with invasion were decreased upon BRAT1 knockdown (BCAS3, GNA12). In addition, pathways associated with the DDR and cell cycle were affected in BRAT1-depleted cells. Importantly, some GBM studies suggested that Cav-1, one of the most increased proteins in NCH644 BRAT1 KD cells, may inhibit migration and invasion by modulating several pathways like MAPK/ERK pathway, crucial for cell proliferation, differentiation, and survival and also inhibiting the PI3K/AKT pathway, important for cell survival and growth. In addition, Cav-1 is essential for cell adhesion and migration. signalling pathways involved in cytoskeletal dynamics and cell motility (Parat & Riggins, 2012). Nonetheless, Cav-1 appears to act in a highly context-dependent manner and has been suggested to enhance migration in other cancer types (Núñez-Wehinger *et al*, 2014). Notably, our global proteomic and phospho-proteomic analysis supports the notion that BRAT1 influences the stemness capacity in NCH644 (ALDH1, PROM1, SOX11, NANOG, RBM15B) and is linked to the DDR through ATM and ATR kinases (Adamski *et al*, 2017; Wang *et al*, 2021). This observation being probably related to the resistance to radiotherapy (Sheehan *et al*, 2010; Arceci, 2008). Further, (Liu *et al*, 2020) explored ATM’s involvement in transforming primary human cells into cancer stem cells. Additionally, kinases linked to migration are suppressed in BRAT1 KD cells including MAPK3 (ERK1) and MAPK8 (JNK1), two members of the MAPK signalling pathway regulating both cell migration and invasion (Huang *et al*, 2004). Additionally, EIF4G1, an EIF4G complex component, that is involved in translation initiation, interacts with MAPK pathway components and was shown to impact mTOR signaling and other downstream signaling cascades in head and neck cancer, was also found to be attenuated after BRAT1 KD. Interestingly, inhibition of the EIF4G complex impaires clonogenicity, tumor sphere formation and cell invasion, with higher EIF4G1 mRNA levels associated with an impaired patient survival in various tumor types (Martínez-Montemayor *et al*, 2010; Tan *et al*, 2019; Jaiswal *et al*, 2019). CDK5 and PAK1 that were also suppressed in BRAT1 KD cells, were previously implicated in cytoskeletal dynamics, cancer cell migration and invasion, suggesting their possible involvement in the pro-migratory role of BRAT1 (Martínez-Montemayor *et al*, 2010; Sharma *et al*, 2002; Bisht *et al*, 2015; Zhou *et al*, 2014). A further candidate, DPYSL2 (CRMP2) is associated with cytoskeletal dynamics, cancer cell migration, invasion, modulation of Rho GTPases and mTOR signaling and promotes axon formation by transporting tubulin heterodimers and oligomers to the tip of the growing axon, as well as by promoting the assembly of tubulin subunits onto the ends of existing microtubules (Pham *et al*, 2016; Conde & Cáceres, 2009). These candidate proteins suggest that BRAT1 is required for migration.

To asses therapeutic implications of the results, we used CurD to target the hitherto “undruggable” protein BRAT1 for proteasomal degradation (Cui *et al*, 2021). Similar, to the findings obtained with BRAT1 KD, we observed anti-migratory effects of CurD treatment, and decreased DDR efficiency after IR. Since the bioavailability of CurD and its ability of passing the blood-brain-barrier has not yet been elucidated yet, we used an OTC transplantation model as a translational *ex vivo* assay and combined it with LSFM to gain insights into the effects of CurD in mouse brain tissue on a single cell level (Haydo *et al*, 2023). CurD enhanced the effect of IR on tumor growth, and OTCxLSFM studies revealed that the pro-migratory phenotype of BRAT1-proficient tumors was curtailed by CurD.

In summary, we find that BRAT1 deficiency causes partial failure of DDR and prevents GBM cells from invasion and migration. Results of knockdown are recapitulated with curcusone D as a pharmacologic tool, hence suggesting that targeting BRAT1 might be a reasonable approach to reduce radioresistance of GBM and limit invasiveness.

## Conclusion

Our study highlights the significance of BRAT1 as a critical factor driving the aggressive pro-migratory and pro-invasive phenotype of GBM. By elucidating its role in enhancing DNA damage response, tumor migration, and invasion, we identified BRAT1 as a putative therapeutic target for GBM. Our findings demonstrate that depletion of BRAT1 sensitizes GBM cells to radiation treatment and inhibits tumor growth and invasion in *in vitro* and *in vivo* models. Furthermore, the use of CurD significantly reduces tumor migration and invasion in GSCs. In combination with radiation treatment, CurD enhanced IR-mediated inhibition of tumor growth and reduced tumor infiltration into brain tissue. Thus, targeting BRAT1 with CurD is a putative novel strategy for GBM therapy.

## 4 Materials and Methods

### 4.1 Analysis of public datasets

Analysis of the Gravendeel dataset (Gravendeel *et al*, 2009) was performed and plots were exported via the GlioVis portal (http://gliovis.bioinfo.cnio.es/) (Bowman *et al*, 2017). Differences of BRAT1 expression were compared between GBMs and healthy tissues based on histology and the Kaplan-Meier curve was derived from the Gravendeel datasets of all GBM subtypes combined with a median cut-off together with confidence intervals. The CGGA (Chinese Glioma Genome Atlas) dataset (Zhao *et al*, 2017) was analyzed for overall BRAT1 mRNA expression in grade 2 tumors in comparison with tumor grades 3 and 4.

### 4.2 Cells and cell culture

Experiments were performed with NCH644, GS-5 human GSCs (Campos *et al*, 2010) and U251 MG glioblastoma cells (Vaheri *et al*, 1976). NCH644 and GS-5 cells were cultured in Neurobasal medium (Gibco, Darmstadt, Germany) supplemented with 1x B27 supplement, 100 U/mL Penicillin, 100 µg/mL Streptomycin, 1x GlutaMAX (all from Gibco), 20 ng/mL epidermal growth factor, 20 ng/mL fibroblast growth factor (both from Peprotech, Hamburg, Germany) and 2 µg/mL Puromycin dihydrochloride (sc-108071B, Lot # L2214, Santa Cruz Biotechnology, Inc., Heidelberg, Germany). U251 cells were cultured in Dulbecco’s modified Eagle’s medium with GlutaMAX and heat-inactivated 10% fetal calf serum, 100 U/mL Penicillin, 100 µg/mL Streptomycin (all from Gibco) and 1 µg/mL Puromycin dihydrochloride (sc-108071B, Lot # L2214, Santa Cruz Biotechnology, Inc.). Both cell lines were cultured at 37°C and 5% CO_2_ and were passaged twice a week in a ratio of 1:10 or 1:20, respectively.

#### Establishment of stable shRNA BRAT1 knockdown cells and BRAT1 overexpressing cells

Stable shRNA-mediated shBRAT1 cells (pLKO.1-puro_shBRAT1_A (N0000172594), pLKO.1-puro_shBRAT1_C (N0000172960) MISSION® (Sigma-Aldrich, Darmstadt, Germany) or control cells expressing non-mammalian targeting control shRNA (pLKO.1-Puro_shCtrl; MISSION® SHC002, Sigma-Aldrich) were generated using HEK293T cells. In addition, NCH644 and U251 cells were transduced with V-lentiCMV-Puro for empty vector control cells and V-lentiCMV-hBRAT1-Puro (self-generated) for producing BRAT1 OE cells. Briefly, 150,000 of HEK293T cells were seeded per well into 6-well plates the day before transfection with 2 µg plasmid DNA (pLKO.1-puro), 1.5 µg gag/pol plasmid (psPAX2, addgene #12260) and 0.5 µg VSV-G envelope plasmid (pMD2.G, addgene #12259) in 57 µL Opti-MEM (Invitrogen, Frankfurt am Main, Germany) and 6 µL FuGENE HD (Promega, Walldorf, Germany) transfection reagent. Six hours later, medium was changed, and the viral supernatant was collected after additional 16h and 40h, pooled and filtered through a 0.45 µm filter, followed by dilution of the supernatant with fresh medium in a ratio of 1:1 and addition of 3 µg/mL hexadimethrine bromide (polybrene, Sigma-Aldrich). 5 µg/mL Puromycin were added and maintained to select for positively transduced cells. psPAX2 and pMD2.G were gifts from Didier Trono (Addgene plasmid # 12260; http://n2t.net/addgene:12260; RRID:Addgene_12260; Addgene plasmid # 12259; http://n2t.net/addgene:12259; RRID:Addgene_12259).

### 4.3 Compounds

The BRAT1 inhibitor curcusone D was dissolved and diluted in DMSO (Sigma) and stored at - 20°C. An intermediate dilution of 3 mM was used for all other dilutions. It was prepared and characterized by the laboratory of Mingji Dai (Cui *et al*, 2021).

### 4.4 Irradiation and Antibodies

Cells were irradiated with single doses of 2, 8 or 10 Gy using a linear accelerator (Elekta Synergy, Elekta) with 6 MV photon energy, 100 cm focus to isocenter distance and a dose rate of 6 Gy/min while non-irradiated control cells were kept in a transportation box in parallel, providing the same experimental conditions.

The following primary antibodies and dilutions were used: mouse-anti-γH2AXSer139 1:1,000 (#05-636, clone JBW301, Merck Millipore) and rabbit-anti-53BP1 1:1,000 (NB#100-304, Novus Biologicals) for immunofluorescence; rabbit-anti-BRAT1 1:250 (HPA029455 Sigma) and mouse-anti-GAPDH 1:1,000 (T6199, Sigma-Aldrich) for western blot. For flow cytometry the FITC conjugated primary antibody γH2AXSer139 –FITC 1:250 (16-202A, Merck Millipore) was used.

The following secondary antibodies with corresponding dilutions were used: F(ab’)2-goat-anti-rabbit IgG (H + L) cross-adsorbed secondary antibody; Alexa Fluor 488 1:500 (A-11070, Thermo Fisher); F(ab’)2-goat-anti-mouse IgG (H + L) cross-adsorbed secondary antibody; Alexa Fluor 594 1:500 (A-11020, Thermo Fisher) for immunofluorescence. IRDye 800CW goat-anti-rabbit 1:10,000 (926-32211) and IRDye 680RD goat-anti-mouse 1:10,000 (926-68070; both LI-COR Biosciences) for western blot.

### 4.5 SDS PAGE and Western Blotting

Cell lysis, SDS-PAGE and Western Blotting were performed as previously described (Antonietti *et al*, 2017). In short, membranes were blocked for 1h at room temperature with 5% BSA/TBS-Tween-20 (TBS-T), followed by an incubation with the primary antibodies as detailed above diluted in 5%BSA/TBS-T at 4°C overnight. The secondary antibodies, diluted in 5% BSA/TBS-T, were incubated for 1h at room temperature and signals were detected using a LI-COR Odyssey reader (LI-COR Biosciences).

### 4.6 γH2AX intensity using Flow Cytometry

U251 and NCH644 cells (6 × 10^5^ cells/mL, 0.5 mL/well) were seeded in 24-well plates. The next day, the cells were treated with 3 µM CurD for 48 h and irradiated with 8 Gy (NCH644 GSC) and 10 Gy (U251 GBM cells). Prior to the measurement, the cells were harvested and stained using γH2AXSer139 –FITC 1:250 (16-202A, Merck Millipore). Cells were analyzed with BD Accuri C6 flow-cytometer (BD Biosciences) and data processing was done using BD Accuri C6 software (BD Biosciences) extracting the mean fluorescence intensity (MFI).

### 4.7 Microscopy

#### γH2AX and 53BP1 Immunofluorescence Microscopy

U251 or NCH644 cells per well were seeded on 8-well chamber slides at a density of 8,000 cells/well (Falcon, Corning). For NCH644 cells, the slides were coated with 10 µg/mL laminin (Sigma, L2020) at 4°C overnight before cell seeding. One day after seeding, the cells were irradiated as indicated. After different time points, they were fixed using 4% formaldehyde/0.25% Triton X-100/ PBS, primary antibodies were diluted in 5% BSA/PBS and incubated at 4°C overnight. Secondary antibodies diluted in 5% BSA/PBS were incubated for 1h at room temperature in the dark. Next, the cells were either mounted on ProLong Gold antifade reagent with DAPI (Invitrogen) or Fluoroshield with DAPI histology mounting medium (Sigma). Slides were stored at 4°C until analysis using a Nikon Eclipse TE2000-S inverted fluorescence microscope operated by the software NIS Elements AR version 4.2 (both Nikon Instruments Europe B.V.) at 60x magnification. For the foci assay, at 1 h after IR γH2AX- or 53BP1-positive foci were counted in 15 nuclei per condition in non-IR cells and 24 h after IR foci were counted in 50 cells per condition.

#### Light-Sheet Microscopy

Light-sheet-fluorescence-microscopy (LSFM) was performed by established protocols by (Haydo *et al*, 2023) to analyze the migration of NCH644^GFP^ DMSO/3 µM CurD treated cells. Briefly, LSFM is a fast 3D fluorescence microscopy technique. It uses a laser light sheet to selectively excite fluorescence in one plane at a time, reducing photobleaching and phototoxicity. This allows for rapid imaging of large samples with high spatial and temporal resolution. For LSFM imaging, a custom-built “monolithic digitally-scanned light sheet microscope” (mDSLM) was used (Pampaloni *et al*, 2013). The mDSLM features a motorized xyzϑ-stage placed below the specimen chamber. Light sheet imaging was performed with an Epiplan-Neofluar 2.5×/0.06 illumination objective (Carl Zeiss), an N-Achroplan 10×/0.3 detection objective (Carl Zeiss), and a Neo CCD camera (ANDOR Technology).The analysis procedures as described in (Haydo *et al*, 2023), provides detailed three-dimensional information at the single-cell level, enabling insights into the morphological characteristics of the tumor cells as well as infiltration or migration, can be gained.

### 4.8 Cell-based assays

#### Sphere Formation

This assay was performed following the established protocol of (Haydo *et al*, 2023; Roth *et al*, 2024). Briefly, to assess changes in sphere area/number and stemness after BRAT1 depletion, cells were seeded at 500 cells/well in a 96-well plate, with 5-10 replicates per condition. After incubation at 37°C for 7 days, images were captured using a Tecan Spark plate reader (Tecan) and analyzed with Fiji ImageJ software (Schindelin *et al*, 2012) to determine mean sphere area and count.

#### Wound Healing Assay – Cell Migration Assay

To investigate the impact of the BRAT1 inhibitor CurD or BRAT1-depletion on cell migration, we employed a wound healing assay using IBIDI chambers (ibidi GmbH). IBIDI inserts offer consistent wall width (500 µm), minimizing experimental error compared to traditional scratch assays. U251 GBM cells were seeded in IBIDI chambers a day prior to treatment. Measurements were taken at 0 h, 24 h, 48 h and 72 h post-treatment using a Tecan Spark reader, with gap width was quantified as percentage from baseline using Fiji ImageJ software (Schindelin *et al*, 2012) to assess migration dynamics.

#### Transwell Migration

The transwell migration (cellQART®) assay measures the ability of cells to migrate through a porous membrane towards a chemoattractant. In this study, NCH644 GSCs were seeded (40,000 cells/chamber) in the upper chamber, and those migrating to the lower chamber are quantified after a defined period. Briefly, the transwell chamber consists of an upper chamber (insert) and a lower chamber (well). GSCs were seeded into the upper chamber of the transwell insert and the lower chamber was filled with culture medium with/without 3 µM CurD. The chambers were fixed after 48 h of incubation. Non-migrated cells on the upper surface of the membrane were removed by gentle wiping with a cotton swab and washing. Migrated cells on the lower surface of the membrane were fixed using methanol, stained with crystal violet and quantified using the Nikon Eclipse TE2000-S inverted fluorescence microscope operated by the software NIS Elements AR version 4.2 (both Nikon Instruments Europe B.V.) at 20x magnification and the number of migrated cells was quantified.

#### Sphere Migration

This assay was performed following an established protocol by (Haydo *et al*, 2023). Briefly, to analyze migration in GSC line GS-5 (Günther *et al*, 2008) tumor spheres, 2,000 to 3,000 cells were seeded into U-shaped 96-well plates and after 48 h transferred into conventional 96-well plates previously coated with laminin. Plates were prewarmed, and spheres allowed to adhere for 30 minutes. Images were taken using a Tecan Spark plate reader (Tecan) before and after 3 µM CurD and DMSO control treatment. Migration was observed up to 72 h. Images were captured every 24 h. Fiji software (Schindelin *et al*, 2012) was used to analyze images, measuring migration distance in micrometers from the edge of the spheres.

### 4.9 Organotypic slice cultures and ex vivo tumor growth assay

For generating adult organotypic slice cultures (OTC) and perform *ex vivo* tumor growth assay we followed established procedures by (Haydo *et al*, 2023; Linder *et al*, 2019; Gerstmeier *et al*, 2021). Briefly, this experiment utilizes OTC, by dissecting mouse brains and generating slices into 150µm thick slices using a vibratome. These slices were cultured on Millicell inserts (Sigma) in six-well plates with FCS-free medium. Tumor spheres were placed onto the brain slices, and treatments (5 µM CurD and 2 Gy fractioned IR) were administered every three days (Monday, Wednesday and Friday).

### 4.10 Proteomic and phospho-proteomic sample preparation and analyses

Proteomic and phosphoproteomic sample preparation and data analyses was performed by established protocols (Linder *et al*, 2022). Briefly, the cells were grown until confluence and were lysed and the proteins were extracted. The proteins were digested using Trypsin (Promega; V5113) and LysC (WakoChemicals). After purification using SepPak C18 columns (Waters; WAT054955) the peptides were TMT-labeled (Thermo Fisher Scientific; 90061;TH266884) and fractionated with the High pH Reversed-phase Fractionation Kit (Thermo Fisher Scientific). After HPLC mediated separation the solutions were directly sprayed into a QExactive HF mass spectrometer and the RAW data was processed with Proteome Discoverer 2.2 software (Thermo Fisher Scientific). TMTpro reporter abundances were extracted and used for plotting and statistical analysis. Further analysis was done on Perseus (v.1.6.5.0) and STRING (v. 12.0). The massspectrometry proteomics data were deposited to the ProteomeXchange Consortium via the PRIDE19,20 partner repository with the data set identifier PXD024802.

### 4.11 Bioinformatic analyses

We employed a combination of bioinformatical tools to analyze protein-protein interaction networks and pathway enrichment, providing insights into the functional relationships and regulatory mechanisms underlying complex biological processes, upon BRAT1 depletion. First, the STRING database (Szklarczyk *et al*, 2019) was utilized to explore known and predicted protein-protein interactions, allowing for the construction of comprehensive interaction networks. Additionally, the Kinase Substrate Enrichment Analysis (KSEA) tool https://casecpb.shinyapps.io/ksea/ (Wiredja *et al*, 2017; Horn *et al*, 2014; Hornbeck *et al*, 2012; Casado *et al*, 2013) was utilized to investigate kinase-substrate relationships and predict potential kinase activities regulating cellular signaling pathways in our U251 phospho-proteomic dataset. This approach provided valuable information on kinase-mediated signaling cascades and their implications in cellular physiology and disease. Analyzing the attenuated kinases in NCH644 GCSs, the LFC were displayed, indicating the changes after BRAT1 depletion. In all, by integrating these bioinformatical tools, we were able to comprehensively analyze protein interaction networks, decipher enriched biological pathways and predict kinase activities, ultimately enhancing our understanding of the molecular mechanisms governing BRAT1 depletion.

### 4.12 Animal experiments

Mouse in vivo experiments was conducted to investigate the role of BRAT1 in promoting tumor growth/invasion and its effect on overall survival using an orthotopic GBM mouse model. Athymic nude mice (n = 15) were used due to their lack of an innate immune system, making it possible to transplant human NCH644 shCtrl and NCH644 shBRAT1 GSC cells into the brain. The animals were housed in type II long cages, purchased from Charles River, and were kept in the “Georg-Speyer-Haus” (Frankfurt, Germany) during experiments. They had free access to food and water. All animal procedures were conducted in accordance with a valid approval (FK-1116) by the Regierungspräsidium Darmstadt and adhered to ARRIVE guidelines set out to reduce animal numbers and minimize suffering. Distinct guidelines and pain scores (score sheets) were established to observe the behavior of the mice upon tumor implantation surgery. Mice were weighted two times a week and checked every two days. If scores exceeded the defined cut off scores for pain and well-being, respective mice were euthanized using a CO_2_ chamber followed by cervical dislocation. Throughout the experiment, efforts were made to minimize the number of animals and to refine experimental procedures to reduce potential harm or discomfort.

### Statistics

All statistical analyses applied are either two-tailed, unpaired t-test for comparison of parametric and normally distributed data of two groups. If prerequisites for t-tests were not met, the non-paramtric Mann–Whitney U test was used as alternative. Survival data were analysed using or Wilcoxon signed rank tests. Multiple groups were compared using one-way ANOVA with Tukey’s or Dunnetts’s multiple-comparison or multivariate ANOVA with Tukey’s multiple-comparison test. Statistical analyses were done with GraphPad Prism 9 and 10 (GraphPad Software, La Jolla, CA, USA). The statistical tests are detailed in the respective figure legends.

## Author Contributions

**Alicia Haydo:** Data curation; software; formal analysis; validation; investigation; visualization; methodology; writing – original draft; writing – review and editing. **Jennifer Schmidt:** Data curation; formal analysis; validation; investigation. **Alisha Crider:** Data curation; formal analysis; validation; investigation. **Tim Kögler:** Data curation; formal analysis; validation; investigation. **Johanna Ertl:** Data curation; formal analysis; validation; investigation. **Stephanie Hehlgans:** Data curation; methodology; resources, writing – original draft. **Marina E. Hoffmann:** Data curation; formal analysis; software; methodology. **Rajeshwari Rathore:** Formal analysis. **Ömer Güllülü:** Methodology, validation. **Yecheng Wang:** Resources. **Xiangke Zhang:** Resources. **Christel Herold-Mende:** Resources. **Francesco Pampaloni:** Resources. **Irmgard Tegeder:** Data curation; resources, writing – original draft. **Ivan Dikic:** Resources. **Mingji Dai:** Resources. **Franz Rödel**: Data curation; formal analysis; validation; resources, writing – original draft. **Donat Kögel:** Validation; resources; supervision; funding acquisition; methodology; writing – original draft; project administration; writing – review and editing. **Benedikt Linder:** Conceptualization; resources; supervision; funding acquisition; methodology; writing – original draft; project administration; writing – review and editing.

## Acknowledgments

The authors would like to thank Hildegard König for her ongoing excellent technical assistance and Martina Leskova for her support during the radiation treatments. The authors would like to thank Dr. Stefan Liebner and the Edinger Institute for assistance with the epifluorescence microscopy analysis. Additional thank is directed towards Margareta Kolaric for her outstanding animal care.

## Supplemental Figures

**Supplemental Figure 1.**
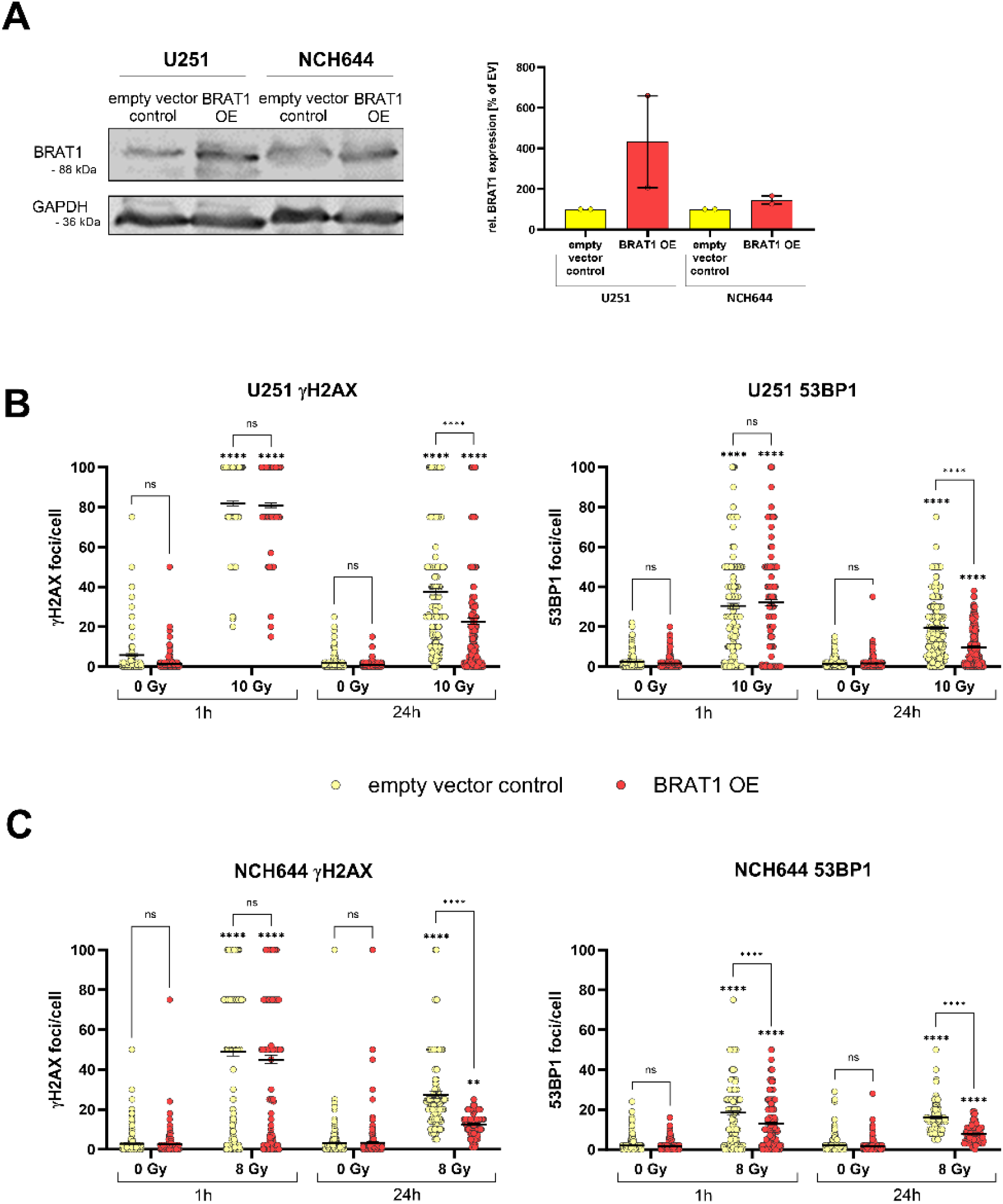
Generating BRAT1 overexpressing cells. (A) Western blot and densitometric validation of stable BRAT1-OE in U251 and NCH644 cells and respective empty vector controls. The quantification shows an increase of BRAT1 expression in the overexpressing cells (BRAT1-OE); GAPDH as housekeeping protein. (B) γH2AX and 53BP1 foci assay of U251 empty vector controls and BRAT1-OE cells. The fluorescence analyses, was performed after 1 h and 24 h of 10 Gy IR. (C) γH2AX and 53BP1 foci assay of NCH644 empty vector controls and BRAT1-OE cells. Analysis as above at 1 h and 24 h of 8 Gy IR. At 1 h after IR γH2AX- or 53BP1-positive foci were counted in 15 cells per condition; in non-IR cells and 24 h after IR, foci in 50 cells were counted per condition. The experiment was performed 3 times in biological replicates and results were pooled with overall n≥15. Statistics: Two-way ANOVA with Tukey’s multiple comparisons test. Error bars are SEM. ns = not significant; ** p < 0.01; **** p < 0.0001 against respective non-IR cells or as indicated.

**Supplemental Figure 2.**
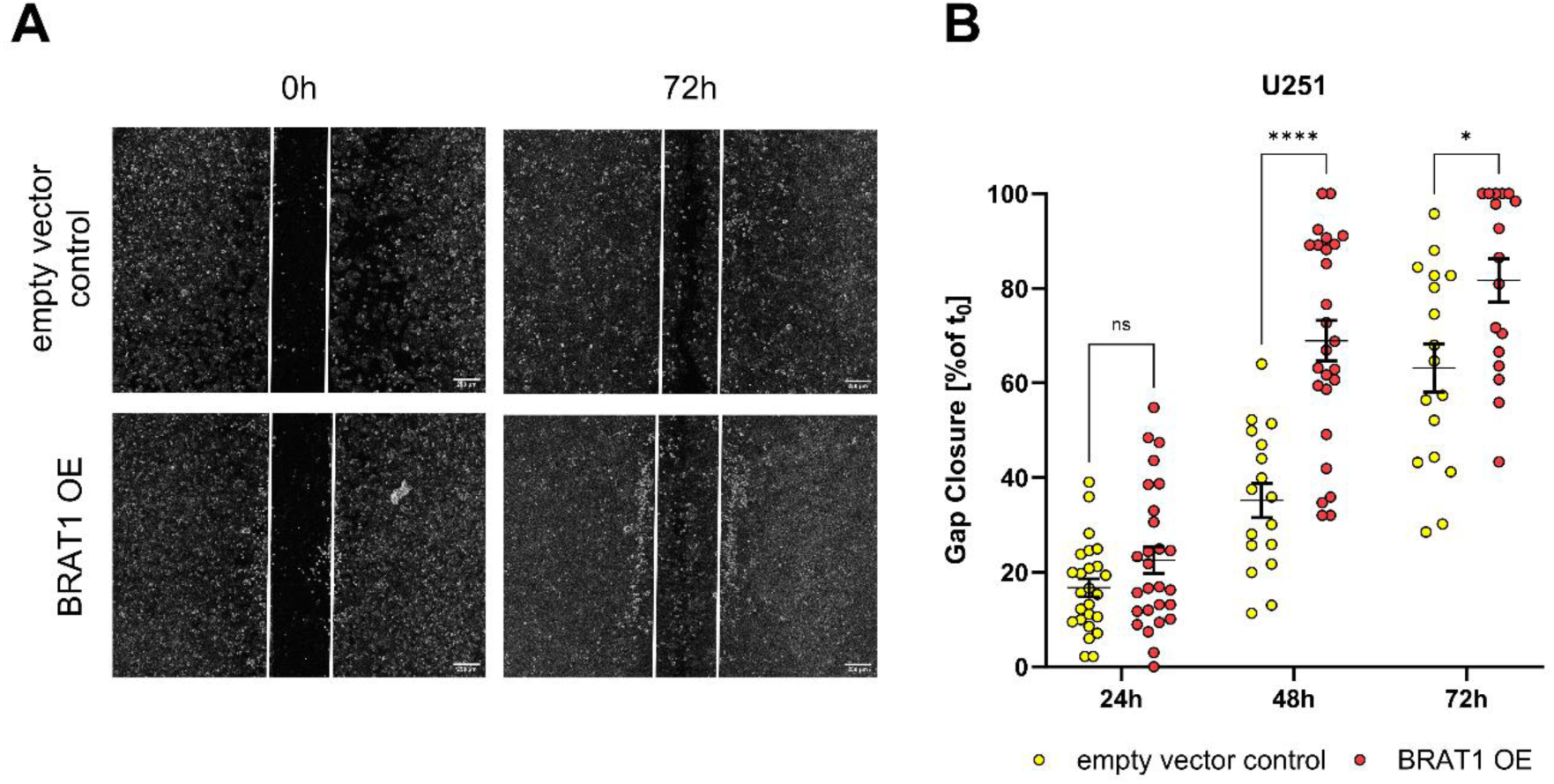
BRAT1 overexpression increases migration capacity. (A) IBIDI migration assay images show the extent of gap closure after 72 h in U251 shCtrl and BRAT1-OE cells; scale bar = 200 µm. (B) Quantification of IBIDI migration assay at 24 h, 48 h and 72 h, presented as gap closure in percentage of baseline at 0 h. The experiment was performed 3 times in biological replicates. Statistics: One-way ANOVA with Tukey’s multiple comparisons test. Error bars are SEM. ns = not significant; * p < 0.05; **** p < 0.0001 against respective shCtrl or as indicated.

**Supplemental Figure 3.**
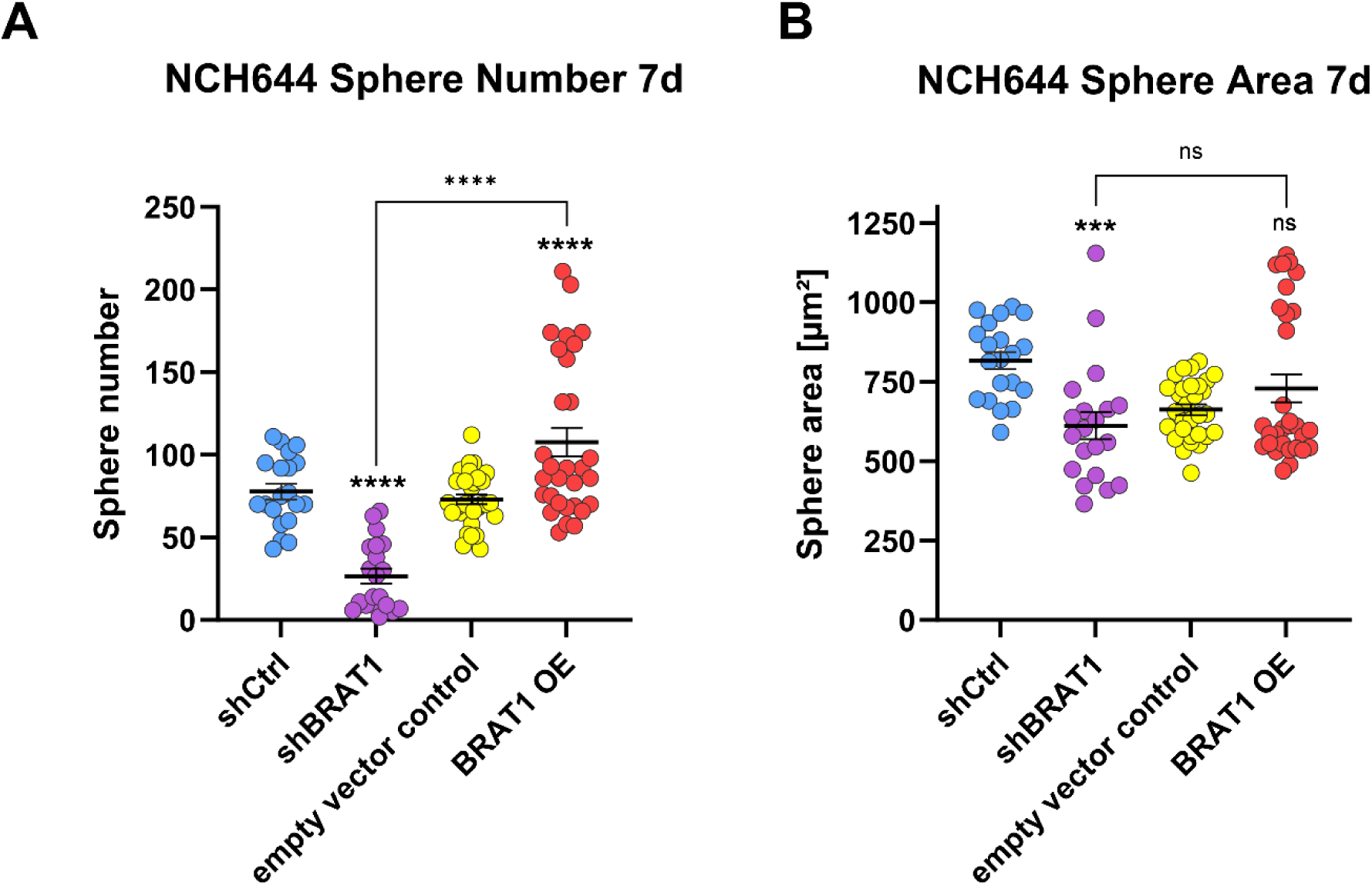
BRAT1 influences stemness capacity to an extent. Sphere numbers and sizes were compared in GSC line NCH644 shCtrl and NCH644 shBRAT1 and NCH644 empty vector control and NCH644 BRAT1-OE cells. (A) Sphere number of NCH644 cells after 7 d of incubation. (B) Sphere area of NCH644 cells after 7 d of incubation. The experiment was performed 3 times in biological replicates, with pooled data resulted in n ≥ 5. Statistics: Two-Way ANOVA with Tukey’s multiple comparisons test. Error bars are SEM. ns = not significant; *** p < 0.001; **** p < 0.0001 against respective control cells (shCtrl for shBRAT1 and empty vector control for BRAT1-OE) or as indicated.

**Supplemental Figure 4.**
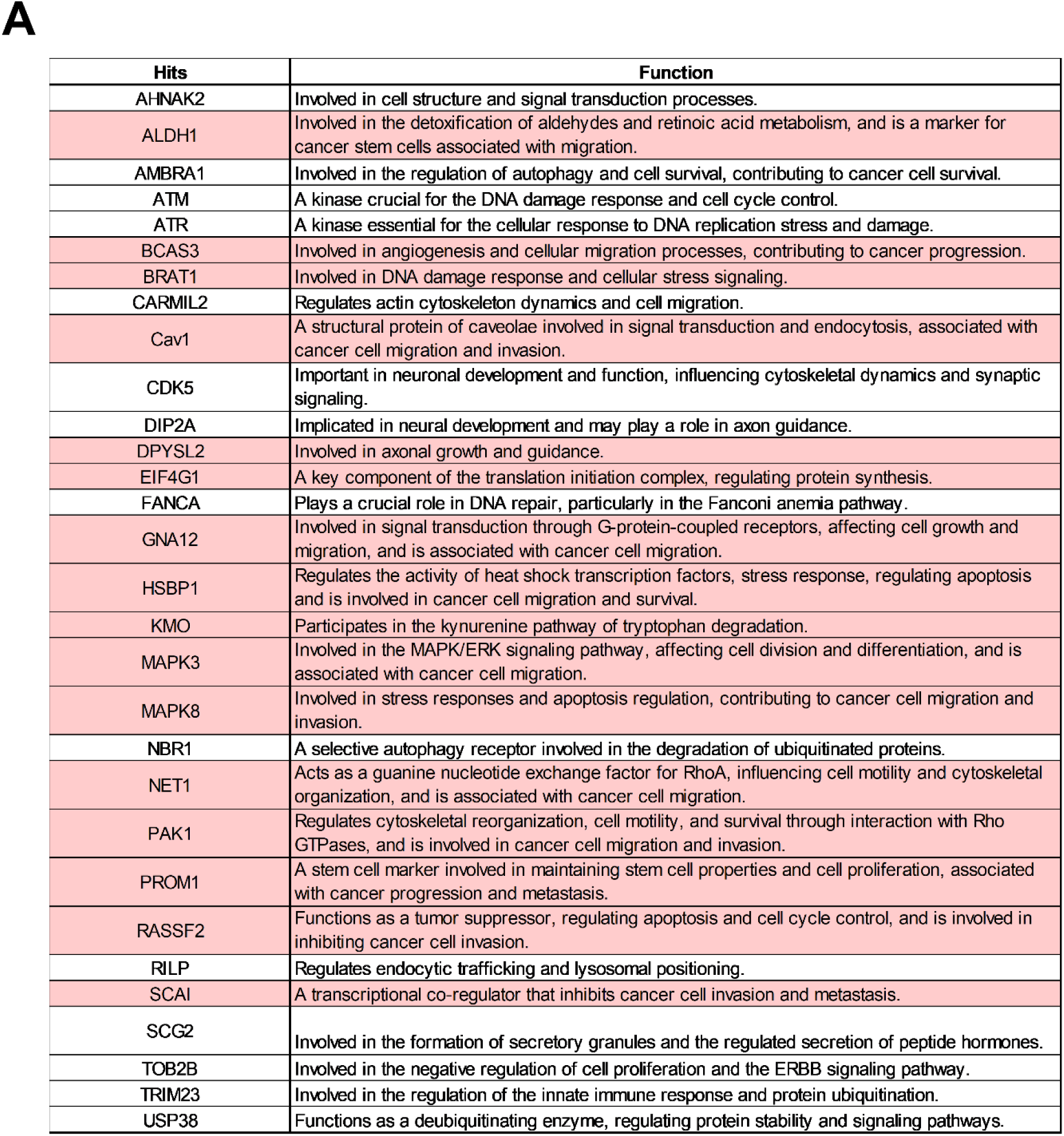
Selected protein hits and their main function. Ordered alphabetically and as displayed in figure 4 and mentioned in the text. “Movement” associated hits are highlighted in light-red.

**Supplemental Figure 5.**
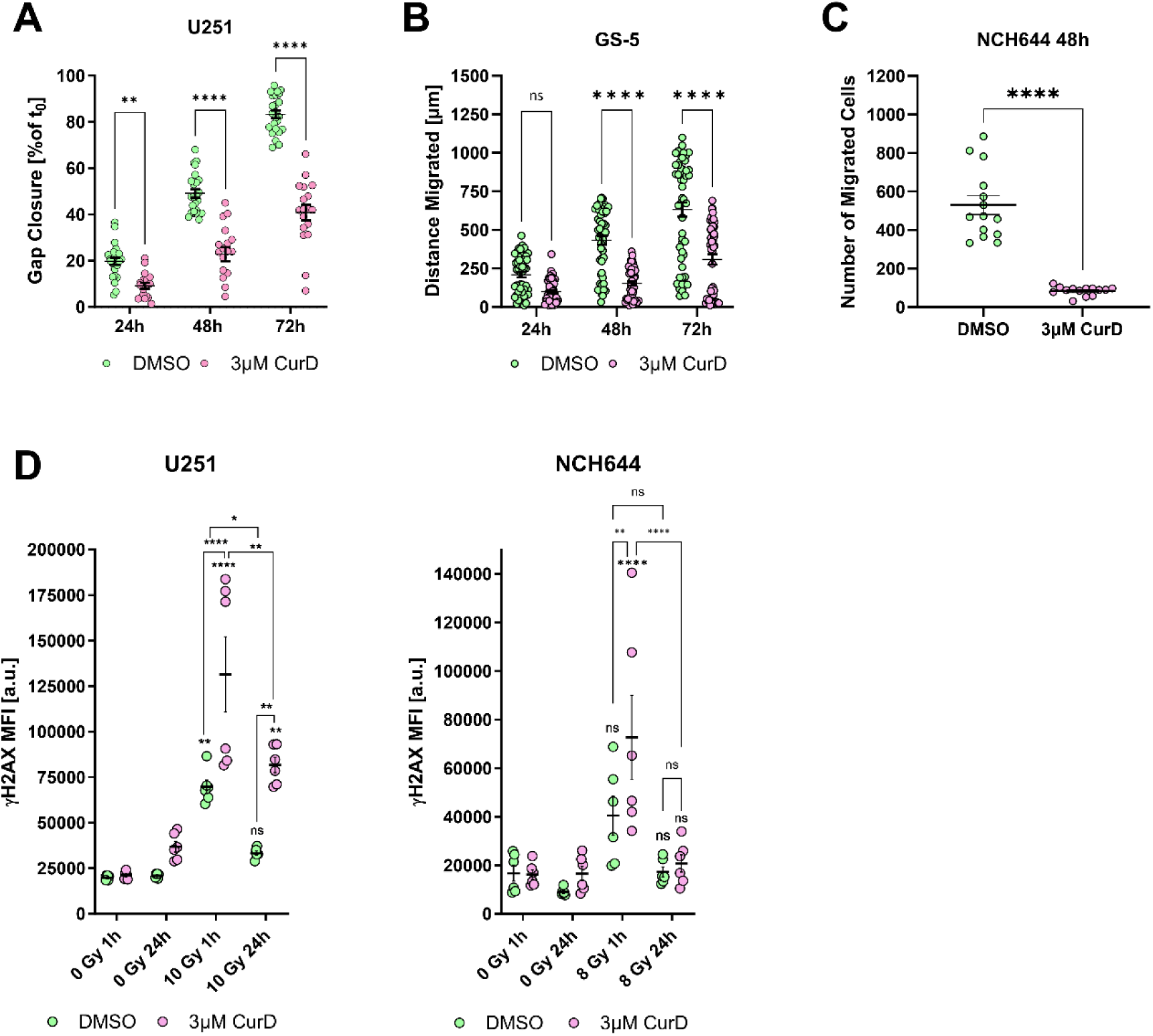
Curcusone D influences migration and leads to a delayed DNA damage response and synergizes with IR-treatment in GBM and GSC lines. All cells were treated with 3 µM CurD or DMSO as solvent control. (A) U251 GBM quantification of IBIDI migration assay displaying 24 h, 48 h and 72 h of gap closure in %. The experiment was performed 3 times in biological replicates with overall n ≥ 3. (B) Sphere migration assay of GSC line GS-5 at 24 h, 48 h and 72 h presented as distance of migrated cells in µm. The experiment was performed 3 times in biological replicates with overall n ≥ 3. (C) Transwell migration of NCH644 cells at 48 h shows number of migrated cells. The experiment was performed 3 times in biological replicates with overall n ≥ 3. (D) γH2AX mean fluorescence intensity (MFI) measured via flow cytometry in U251 and NCH644. In addition to drug treatment, cells were irradiated with 10 Gy for U251 and 8 Gy for NCH644 cells. The experiment was performed 3 times in biological replicates with overall n ≥ 3. Statistics: A-C: Unpaired Mann–Whitney test. D: Two-way ANOVA with Tukey’s multiple comparisons test. Error bars are SEM, ns = not significant; * p < 0.05; ** p < 0.01; **** p < 0.0001 against respective DMSO treated cells or as indicated.

